# Caspar, an adapter for VAP and TER94 delays progression of disease by regulating glial inflammation in a *Drosophila* model of ALS8

**DOI:** 10.1101/2021.04.07.438776

**Authors:** Shweta Tendulkar, Sushmitha Hegde, Aparna Thulasidharan, Lovleen Garg, Bhagyashree Kaduskar, Anuradha Ratnaparkhi, Girish S Ratnaparkhi

**Affiliations:** Indian Institute of Science Education & Research (IISER) Pune 411008, INDIA; Agharkar Research Institute, Pune 411004, INDIA

**Keywords:** Amyotrophic Lateral Sclerosis, Proteostasis, FAF1, NFkappa B

## Abstract

Amyotrophic Lateral Sclerosis (ALS) is a fatal, late onset, progressive motor neurodegenerative disorder. We have been studying cellular and molecular mechanisms involved in ALS using a vesicle-associated membrane protein-associated protein B (VAPB/ALS8) *Drosophila* model, which mimics many systemic aspects of the human disease. Here, we show that the ER resident VAPB interacts with Caspar, an ortholog of human fas associated factor 1 (FAF1). Caspar, in turn, interacts with transitional endoplasmic reticulum ATPase (TER94), a fly ortholog of ALS14 (VCP/p97, Valosin-containing protein), via its UBX domain and poly-ubiqutinated proteins with its UBA domain. Caspar overexpression in the glia extends lifespan and also slows the progression of motor dysfunction in the ALS8 model, a phenomenon that we ascribe to its ability to restrain age-dependant inflammation, modulated by Relish/NFκB signalling.

We hypothesize that Caspar is a key molecule in the pathogenesis of ALS. Caspar connects the plasma membrane (PM) localized immune signalosome to the ER based VAPB degradative machinery, presumably at PM:ER contact sites. The Caspar:TER94:VAPB complex appears to be a strong candidate for regulating both protein homeostasis and NFκB signalling. These, in turn, regulate glial inflammation and determine progression of disease. Our study projects human FAF1 as an important protein target to alleviate the progression of motor neuron disease.

## INTRODUCTION

Amyotrophic Lateral Sclerosis (ALS) is a human neurodegenerative disease that leads to the death of motor neurons. This disorder usually affects individuals late in life, with 90-95% cases being ‘sporadic’, indicating that the disease is not inherited. In about 5-10% of cases the disease is ‘familial’, where the disease is inherited across generations (Cleveland and Rothstein 2001; Pasinelli and Brown 2006; Mitchell and Borasio 2007). Despite the discovery of SOD1 as the first familial locus for ALS nearly three decades ago in 1993 (Deng *et al*. 1993; Rosen *et al*. 1993), and the large volume of work, an understanding of the mechanistic aspects of disease initiation and progression has been elusive. One major reason for the delay has been the identification of more than two dozen ALS causative loci in humans, with each locus being distinct and unrelated to each other in terms of their physiological function (Abel *et al*. 2012; Robberecht and Philips 2013). For example, SOD1/ALS1 is primarily a regulator of cytoplasmic Reactive Oxygen Species (ROS), Senataxin (SETX)/ALS4 (Chance *et al*. 1998) works as a RNA or DNA helicase, FUS/ALS6 (Vance *et al*. 2009) is a transcriptional activator, VAPB/ALS8 (Nishimura *et al*. 2004) is an ER membrane resident single pass protein that helps maintain intracellular membrane contact sites, while KIFA/ALS25 (Nicolas *et al*. 2018) is a microtubule based motor protein. Nonetheless, specific mutations in each of these genes leads to development of the disease. Whether the pathway leading to motor neuron death is distinct for each locus or is caused by convergence to a common downstream cellular mechanism is not completely clear. A universal mechanistic model for the disease should be able to explain the convergence to the common end point, by incorporating a related set of cellular pathways that are perturbed by each of the causative loci. This would presuppose that subsets of ALS causing genes are connected, being part of genetic sub-network(s) in the cell, regulating common or overlapping physiological functions. If true, then it should be possible to uncover genetic relationships between different ALS loci and link these interactions to a set of common cellular pathways. This is best done in animal models where genetic tools to elucidate such interactions exist. In the past decade many such interactions have been discovered and cellular themes underscored. For example, multiple ALS loci have been found to be involved in functions related to RNA metabolism. These include *TDP-43, FUS, ATXN2, TAF15, EWSR1, hnRNPA1, hnRNPA2/B1, MATR3* and *TIA1*. Another common theme is proteostasis, involving both production of proteins - *TARDBP, FUS, TAF15* as well as degradation-*VCP, SQSTM1, UBQLN2, OPTN, and CCNF*. Finally, vesicular transport, another potential fault line for ALS, involves *C9ORF72, VAPB, OPTN, CHMP2B, PFN1, TUBA4A, ANXA11, ATXN2, NEFH, PRPH, SPG11, SIGMAR1, GRN, DCTN1, KIF5A and ALSIN*. Further support for the genetic sub-network hypothesis is the finding that ALS is in all possibility a polygenic disease with several genes contributing to initiation and progression in both sporadic (sALS) and familial (fALS) cases (Van Blitterswijk *et al*. 2012; Leblond *et al*. 2014; Renton *et al*. 2014).

VAPB was the 8^th^ fALS locus to be discovered (Nishimura *et al*. 2004) and has been used to model the disease in both invertebrates (Pennetta *et al*. 2002; Tsuda *et al*. 2008; Han *et al*. 2012; Moustaqim-Barrette *et al*. 2014) and vertebrates (Teuling *et al*. 2007; Larroquette *et al*. 2015). VAPB is a ubiquitous, tail anchored ER protein, enriched in contact sites where ER is juxtaposed with other intracellular membranes (reviewed by (Lev *et al*. 2008; Murphy and Levine 2016; Kamemura and Chihara 2019)). VAPB associates with a number of partner proteins, re-enforcing membrane-membrane distance and architecture (Kamemura and Chihara 2019) and also assisting in roles performed by its numerous interacting partners (Murphy and Levine 2016; Kamemura and Chihara 2019). Recent studies have focused on elucidation of specific roles for VAPB with each individual partner protein (Lindhout *et al*. 2019; Mao *et al*. 2019; Cabukusta *et al*. 2020; Di Mattia *et al*. 2020), broadening our understanding of the workings of the VAPB network.

We have previously sought to uncover the gene regulatory network for the *Drosophila* ortholog of human VAPB, VAP33A/CG5014 (VAP here onwards) and to understand genetic relationships between ALS loci through an enhancer-suppressor screen (Deivasigamani *et al*. 2014). The study identified ALS orthologous loci (*SOD1, Alsin, TDP43*) as part of the VAP genetic network and also suggested an interaction with members of the Target of Rapamycin (TOR) signalling pathway (Deivasigamani *et al*. 2014). We subsequently explored the genetic relationship between *VAP* and *SOD1* and found that SOD1 regulation of cellular reactive oxygen species (ROS) levels led to a modulation of VAP^P58S^ aggregates (Chaplot *et al*. 2019), by triggering clearance of cellular inclusions via the ubiquitin-proteasomal system (UPS). In the current study, we explore the relationship between VAP and Transitional endoplasmic reticulum (ER) ATPase (TER94/ALS14; or p97, also called valosin containing protein, VCP). The decision to explore this relationship in detail was based on a targeted, tissue specific genetic screen, described here, carried out in muscles, glia and motor neurons.

In our study, we find that Caspar, an ortholog of human Fas associated factor 1 (FAF1)(Chu *et al*. 1995; Ryu *et al*. 2003) is a physical interactor of both VAP and TER94 (Kang and Yang 2011; Kim *et al*. 2011; Baron *et al*. 2014). We show that Caspar acts as a protein bridge, physically connecting VAP and TER94, that are orthologs of human VAPB/ALS8 and VCP/ALS14 respectively. Interestingly, increased Caspar expression in glia can improve the lifespan of our ALS8 disease model and can also delay the onset of motor dysfunction. The VAP:Caspar:TER94 complex appears to be a component of the ER -associated degradation (ERAD) network, with Caspar regulating the progression of the disease by modulating NFκB signalling. Our findings suggest that Caspar in glia plays a key role in disease progression by regulating age dependant glial inflammation thus highlighting the contribution of non-autonomous players in the development of the disease.

## RESULTS

### A tissue specific enhancer/suppressor screen suggests genetic interactions between *VAP* and *Drosophila* orthologs of ALS loci

In the past we (Ratnaparkhi *et al*. 2008; Deivasigamani *et al*. 2014; Yadav *et al*. 2018; Chaplot *et al*. 2019) and others (Pennetta *et al*. 2002; Chai *et al*. 2008; Tsuda *et al*. 2008; Han *et al*. 2012; Forrest *et al*. 2013; Sanhueza *et al*. 2015) have used VAP^P58S^ overexpression models to understand mechanisms of disease. Here, for our experiments, we chose a ALS8 disease model developed by Hiroshi Tsuda’s laboratory (Moustaqim-Barrette *et al*. 2014), where an endogenous *genomic-VAP* (*gVAP* or *gVAP*^*P58S*^) insert on the third chromosome is used to rescue the lethality of the VAP null (*ΔVAP*). The Tsuda disease model, *ΔVAP;gVAP*^*P58S*^, unlike the control *ΔVAP; gVAP*^*WT*^, has a shorter lifespan, shows progressive motor dysfunction, has VAP^P58S^ inclusions in the brain with cells showing ER expansion and ER stress (Moustaqim-Barrette *et al*. 2014). In our laboratory, we have regenerated the Tsuda model, using the *VAPΔ166* allele instead of the *VAPΔ31* (See Materials & Methods) and have used this for our studies.

In order to study the genetic relationships between VAP and different ALS loci, we chose six *Drosophila* genes, each being an ortholog of a major fALS locus (Fig. 1A). The genes chosen are fly orthologs of human *TARDBP/ALS10, FUS/ALS6, SOD1/ALS1, VCP/ALS14 and SETX/ALS4*, respectively. For each gene, transgenic lines for overexpression (*OE*) and knockdown *(KD)* experiments were procured (See Materials & Methods). The goals of our experiments were twofold: first to test interactions between each gene and VAP using the disease model as a readout. Second, to explore these genetic interactions in the three tissues central to motor neuron disease, namely Neurons, Muscle and Glia. These experiments were driven by the concept that in addition to neuronal cells, non-neuronal cells (Boillee *et al*. 2006) also contribute to the disease. Three drivers were chosen, one for each tissue, *myosin heavy-chain* (*MHC*)-*Gal4* for muscle, *reverse polarity* (*Repo)-Gal4* for glia and *OK6-Gal4* for motor-neuron specific expression.

**Figure 1.**
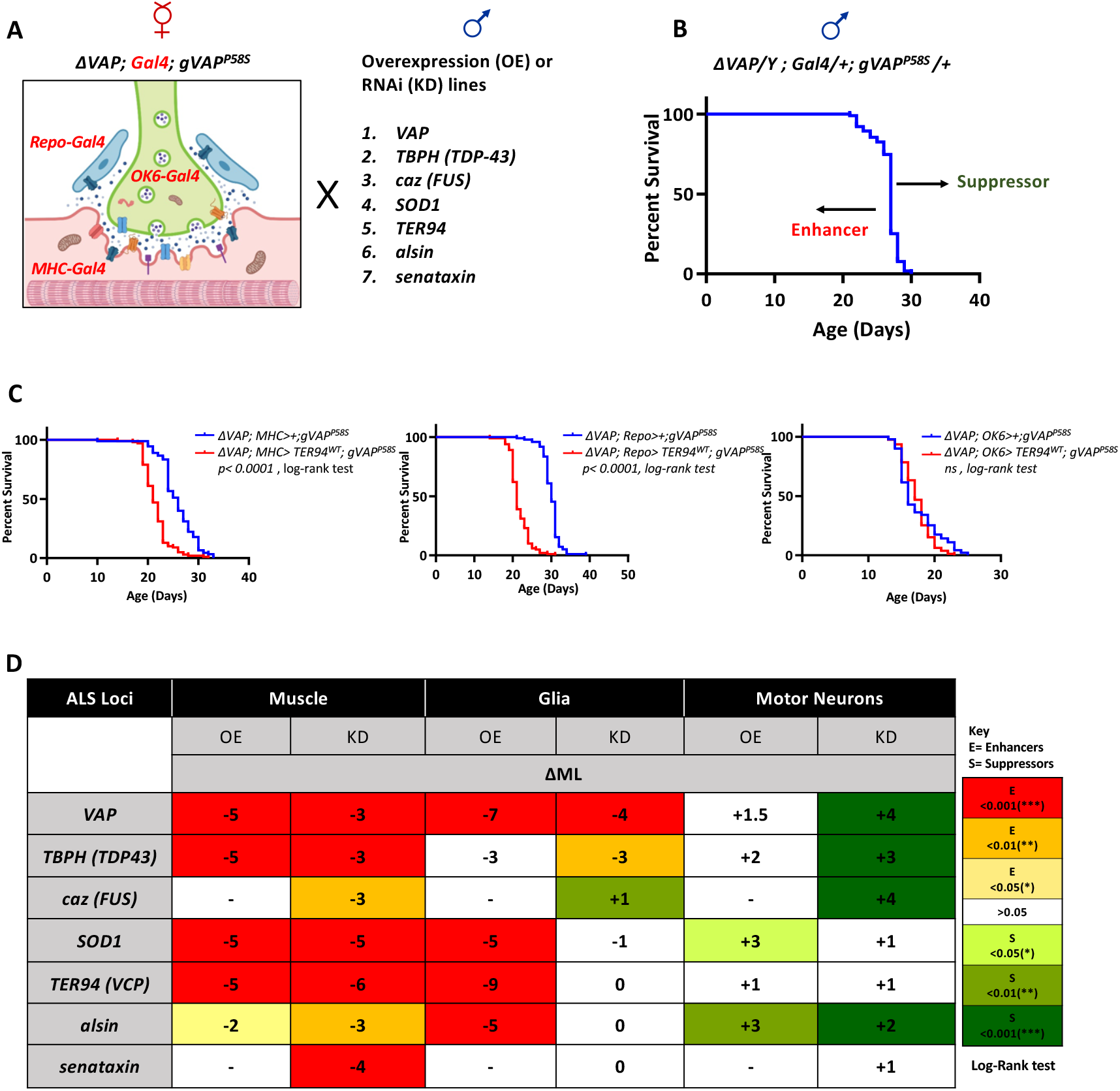
An enhancer/suppressor screen for genetic interaction between ALS orthologous loci in neurons, muscle and glia. **A**. Tissue specific expression in muscle (*∆VAP; MHC-Gal4; gVAP*^*P58S*^), glia (*∆VAP; Repo-Gal4; gVAP*^*P58S*^) and motor neurons (*∆VAP; OK6-Gal4; gVAP*^*P58S*^) was used to overexpress and knockdown *VAP, TBPH, caz, SOD1, TER94, alsin* and *senataxin*. **B**. A schematic for monitoring survival of male animals emerging from the crosses described in panel A. The blue curve represents the control *∆VAP/Y; X-Gal4/+; gVAP*^*P58S*^ /+ fly line i.e. *∆VAP/Y; MHC-Gal4/+; gVAP*^*P58S*^*/+* for the muscle-specific screen, *∆VAP/Y; Repo-Gal4/+; gVAP*^*P58S*^*/+* for the glial-specific screen and *∆VAP/Y; OK6-Gal4/+; gVAP*^*P58S*^*/+* for the motor neuron-specific screen. A shift from the lifespan of the control to the right was defined as a ‘suppression’ of the phenotype while towards the left as ‘enhancement’. The survival analysis was done using the log-rank test survival assay protocol, in Prism 7, over the entire dataset, where each curve was compared to the control and the value of significance was noted. This number is reported (*Suppl. Fig. 1*) and is represented as a colour code in panel D. Additionally, we considered values of ∆ML (Change in Median Lifespan (ML)) = ML (Experiment) – ML (Control) as a simple readout of enhancement or suppression. **C**. Representative graphs for muscle, glial and motor neuronal overexpression of *TER94*. The red curve represents the locus studied (*TER94*). The *p*-value from the log-rank test listed in each graph is the comparison of the red curve with the blue curve (control). The muscle and glial overexpression of *TER94* exhibit enhancement of life span defect (*p< 0*.*0001*; log-rank test) while motor neuronal expression shows no significant change. A complete set of graphs (muscle, glia, neurons) for *KD* and *OE* of all seven genes can be found in *Suppl. Fig. 1*. **D**. Tabular summary of the tissue-specific screen highlights differential interaction of genetic loci with *VAP*^*P58*S^. The numbers in each cell indicate the ∆ML, while the colours indicate different levels of statistical confidence, as per the log-rank test. The ‘red-yellow tones’ mark Enhancers (E) while the ‘green tones’ represent the same for Suppressors (S). ‘-’ in the cell indicates that the experiment was not done. The *caz* overexpression construct is an insert on the X chromosome of the fly line and hence could not be used for male specific assay, while for *senataxin*, the OE line was not available. ‘OE’ stands for overexpression using UAS lines and ‘KD’ stands for knockdown, using RNA interference.

Gal4 drivers for glia, muscle and neurons with insertions in the second chromosome were balanced with *ΔVAP* and *gVAP*^*P58S*^ to generate the following lines: *ΔVAP; Repo-Gal4; gVAP*^*P58S*^, *ΔVAP; OK6-Gal4; gVAP*^*P58S*^, *ΔVAP; MHC-Gal4; gVAP*^*P58S*^ (Fig. 1A). Females of these driver lines were crossed to *OE or KD(RNAi)* lines of *VAP, TBPH, FUS, SOD1, TER94, ALSIN* and *SETX*. Adult males in the F1 generation, which lacked functional *VAP* on the X chromosome (*ΔVAP;* Fig. 1B) were collected and subjected to lifespan analysis as described (Materials & Methods). Survival graphs for each cross are shown (*Suppl. Fig. 1*; Representative graphs in Fig. 1C), with a summary tabulated in Fig. 1D. Statistical analysis using the Log-Rank Test, as detailed in Materials and Methods, were used to compare experimental lifespan curves with controls and change (Δ) in median lifespan (ML) was used as a simple parameter to reflect increase (+) or decrease (-) in lifespan (Fig. 1D).

A summary of results (Fig. 1D) of the differential lifespan analysis (data in *Suppl. Fig 1*), is as follows. In the muscle, both decrease or increase of activity of the genes tested seems to enhance the phenotype with a concomitant decrease of ML. In glia, increased activity of most of the genes (*VAP, SOD1, TER94* and *alsin*) tested, seems to enhance the phenotype with a decrease of ML. In the *KD* studies, *VAP* and *TBPH* lead to decrease in ML while for other genes, the changes in lifespan were not very significant. In neurons, the overall trend pointed to a suppression of the phenotype, with increase in ML for knockdown of *VAP, TBPH, caz and alsin. SOD1* OE, as also *alsin OE* led to a mild suppression of the phenotype. For most other genes, the effect was not significant (*p*>0.05). The strongest enhancement of the lifespan phenotype was for *TER94 OE* with a ΔML ranging from 7-10 days; *TER94 KD*, in contrast, did not influence the lifespan of the *ΔVAP/Y; Repo-Gal4/+; gVAP*^*P58S*^*/+* line significantly (Fig. 1C,D). We chose to further explore, in greater detail, the TER94:VAP interaction, using our disease model. This direction was supported by a study from Gabriela Alexandru’s group (Baron *et al*. 2014), which suggested that VAPB and VCP could interact physically in mammals via FAF1. As a first step, we conducted affinity purification experiments using fly lysates to test if VAP was a physical interactor of Caspar and if so, whether Caspar could be a potential adapter for the VAP:TER94 interaction in our disease model.

### Caspar is a physical interactor of both VAP and TER94

A sequence comparison of Caspar with FAF1 finds that both proteins are of similar size, have the same domain structure (Fig. 2A) and on sequence alignment show 54% sequence similarity overall with ∼34% identity. Like its mammalian ortholog, Caspar contains well characterized, N-terminal, Ubiquitin associated (UBA; SM00165) domain, known to interact with poly-ubiquitinated proteins and a C-terminal ‘Ubiquitin regulatory X’ UBX domain (SM00166), a known VCP/TER94 interactor (Yeung *et al*. 2008). As detailed earlier (Baron *et al*. 2014), the Caspar superfamily is conserved from flies to humans and incorporates a variant of FFAT, ‘two phenylalanines in an acidic tract’ motif, EFFDAxE, (Loewen *et al*. 2003; Loewen and Levine 2005). It is well established that this motif mediates interaction with VAP (Loewen and Levine 2005; Murphy and Levine 2016; Yadav *et al*. 2018). Consistent with this, deletion or modification of this motif led to disruption of the FAF1:VAPB interaction (Baron *et al*. 2014). This strongly suggests that *Drosophila* Caspar, which contains a similar highly conserved FFAT motif (Fig. 2A, lower panel, Underlined) may also be a VAP interactor.

**Figure 2.**
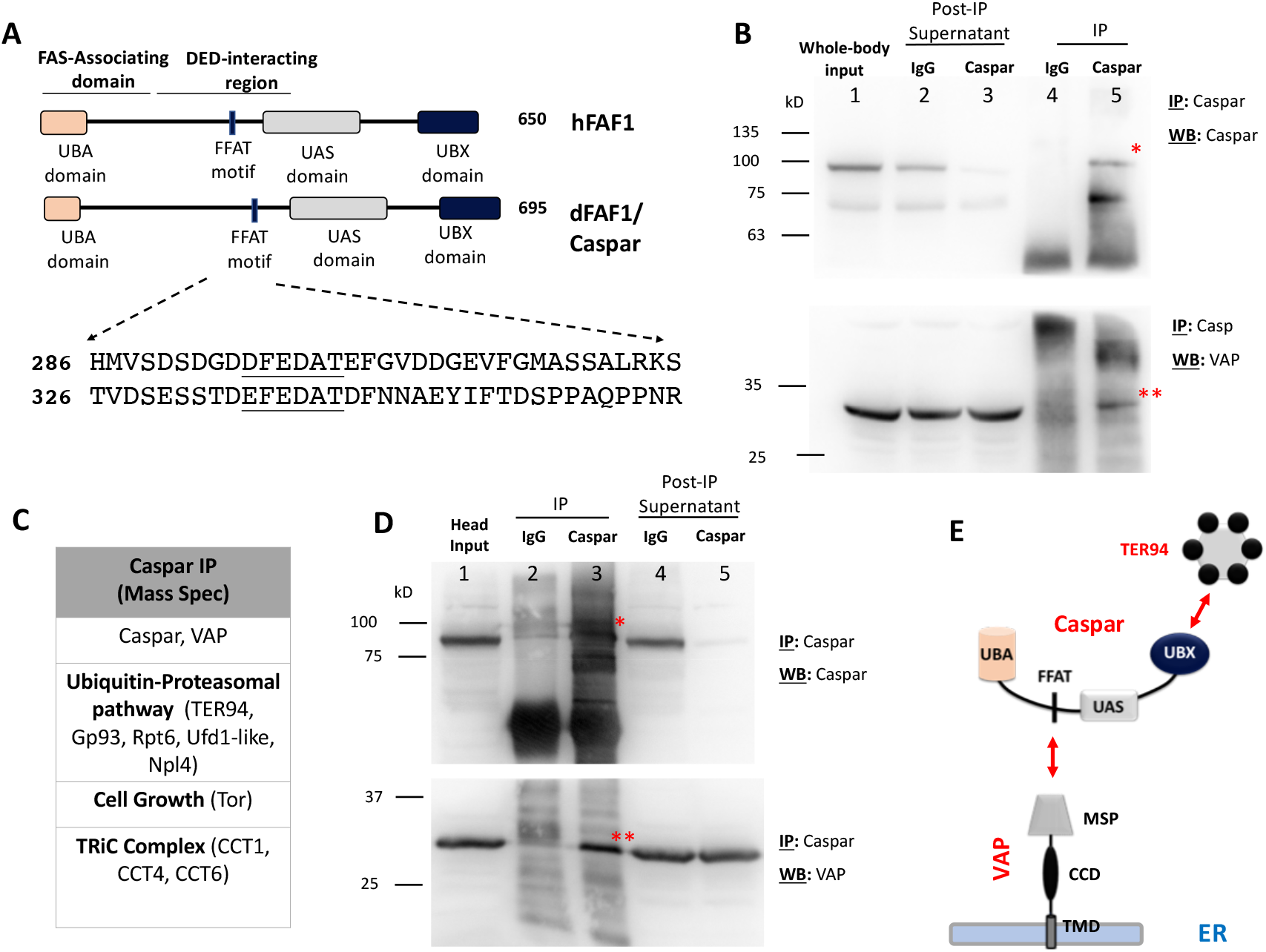
*Drosophila* Caspar is a physical interactor of VAP. **A**. *Drosophila* Caspar is an ortholog of human FAF1. The conserved N-terminal Ub-interacting and C-terminal TER94 interacting domains suggest roles for Caspar as an adapter in the proteasomal degradation. A conserved FFAT motif (amino acids underlined), which is a well characterized VAP interactor is present in both polypeptides. This suggests that FFAT motif in Caspar may be the interface for an interaction with VAP. **B**. An antibody against full length Caspar was generated (Materials and Methods) and validated (*Suppl. Figure 2*). When used for immunoprecipitation (IP) of whole animal lysates, the Rb Caspar antibody could affinity purify Caspar (*, 90 kD, lane 5, top panel) and also VAP (**, 25 kD, lane 5, bottom Panel). A concomitant decrease in Caspar in the supernatant (lane 3, top panel) is also seen, post IP. The ∼70 kD reactive band is a feature of Caspar westerns. **C**. TER94, Gp93, Rpt6, Tor, CCT1 and CCT4 are detected by Mass Spectrometry in the Caspar antibody immune-precipitates, using whole-body and embryonic lysates (*Suppl. Table 1*). Many of these are functionally associated with the ubiquitin-proteasomal system (UPS). **D**. Caspar immune-precipitates using fly head lysates confirm that Caspar (*) is expressed in the head and can be enriched by the Caspar antibody (lane 3, top panel). Also, VAP (**) is a Caspar interactor (lane3, bottom panel). A concomitant decrease in Caspar in the supernatant (lane 5, top panel) is seen, post IP. Further support for the Caspar:VAP interaction comes from a ‘reverse’ IP experiment where an anti-VAP antibody is used for immunoprecipitation and Caspar is enriched (*Suppl. Fig. 2C*). **E**. A model for the interaction of VAP, an ER resident membrane protein, with cytoplasmic Caspar. Mass spectrometry interaction data suggests that TER94 and other proteins of the UPS are part of Caspar protein:protein interaction network in the cell.

To test the VAP:Caspar interaction, we used an anti-Caspar antibody developed in our laboratory, as described in materials & methods, to immuno-precipitate (IP) Caspar and its interactors. The anti-Caspar antibody specifically recognizes a ∼85 kD band in adult whole animal lysates (*Suppl. Fig. 2A*), which is not seen in lysates of homozygous *caspar*^*lof*^ animals (*Suppl. Fig. 2A*). The anti-Caspar antibody IPs could enrich Caspar, as visualized by Western Blotting (Fig. 2B; *, Top panel) and could also pull down VAP (Fig. 2B; **, Bottom panel). To identify other Caspar interactors, we processed the Caspar IPs from adult and embryonic lysates through a Mass Spectrometer and found that Caspar associates with cytoplasmic proteins involved in protein folding/unfolding and ubiquitin mediated degradation (Fig. 2C, *Suppl. Fig. 2D*). In embryonic lysates, which is a rich source of Caspar, VAP is found as the second most enriched protein (after TER94), based on peptide count. In fly head extracts, the anti-Caspar antibody enriches both Caspar (Fig. 2D; *, top panel) and VAP (Fig 2D; **, bottom panel). Further support for the VAP:Caspar interaction comes from a ‘reverse IP’ experiment in fly heads, where a VAP IP could pull down Caspar (*Suppl. Fig. 2C*). The VAP:Caspar physical interaction and the list of associated proteins such as TER94, Ufd1-like and Npl4 gives credence to the idea that Caspar acts as an adapter to bring together VAP and TER94 for an important physiological function, possibly related to protein homeostasis and degradation. Caspar is thus a *bona-fide* interactor of both VAP and TER94, and this protein complex is conserved from flies (VAP:Caspar:TER94) to humans (VAPB:FAF1:VCP; (Baron *et al*. 2014)).

### Expression of Caspar in the glia extends lifespan

The existence of a VAP:Caspar:TER94 complex in flies suggests that TER94 and Caspar may have common or overlapping physiological roles. In order to uncover these we expressed two ALS fly variants of *TER94, TER94*^*A229E*^ and *TER94*^*R152H*^ (Ritson *et al*. 2010; Chang *et al*. 2011) in the glia, muscle and neurons. Expression of *TER94*^*R152H*^ led to a significant extension of lifespan (ΔML=+4.5) when expressed in glia, but not in muscle (ΔML=-2) or neurons (ΔML=+2). *TER94*^*A229E*^ expression in the glia was lethal and adult flies did not emerge (Fig. 3A). R152 is located in the N-terminal CDC48 domain of TER94 and R152H is classified as a dominant active allele (Chang *et al*. 2011; Zhang *et al*. 2015). The N-terminal domain of TER94 is also known to interact with UBX domains, suggesting that the mutation may modulate the strength or dynamics (Zhang *et al*. 2015) of the TER94:Caspar interaction. In order to test the effect of *OE* and *KD* of Caspar in glia, we procured a RNAi line for *caspar* (*UAS-caspar*^*RNAi*^) and also generated *UAS-caspar* lines (See Materials & Methods). *ΔVAP; Repo-Gal4/+; gVAP*^*P58S*^*/UAS-caspar*^*RNAi*^ males had a shorter lifespan (ΔML= -4.0) when compared to the disease model, *ΔVAP/Y; Repo-Gal4/+; gVAP*^*P58S*^*/+* (Fig 3 A, B). Intriguingly, overexpression of *caspar* in the glia (*ΔVAP; Repo-Gal4/+; gVAP*^*P58S*^*/UAS-caspar*), significantly increased lifespan (ΔML=+7.5)(Fig. 3B) of the disease model. In repeat experiments, ΔML ranged from 7-9 days, with the extension of lifespan always greater than the (*ΔVAP; Repo-Gal4/+; gVAP*^*P58S*^*/UAS-TER* ^R152H^) experiment, where the ΔML ranged from 4-6 days. The dramatic increase was made even more significant because overexpression of c*aspar* in a wild-type background (Fig. 3 C,D) severely shortened lifespan (ΔML=-17)(Fig. 3D), while *caspar KD* increased lifespan (ΔML=+4). For *TER94*, both *OE* and *KD* decreased ML, in wild-type animals, by 3.5 and 9 days respectively, pointing to the importance of maintaining homeostatic levels of TER94 in the glia. Thus, the increased lifespan seen after *TER94*^*R152H*^ and *caspar*^*WT*^ glial *OE* strongly suggests an important physiological role for the VAP:Caspar:TER94 complex in our disease model.

**Figure 3.**
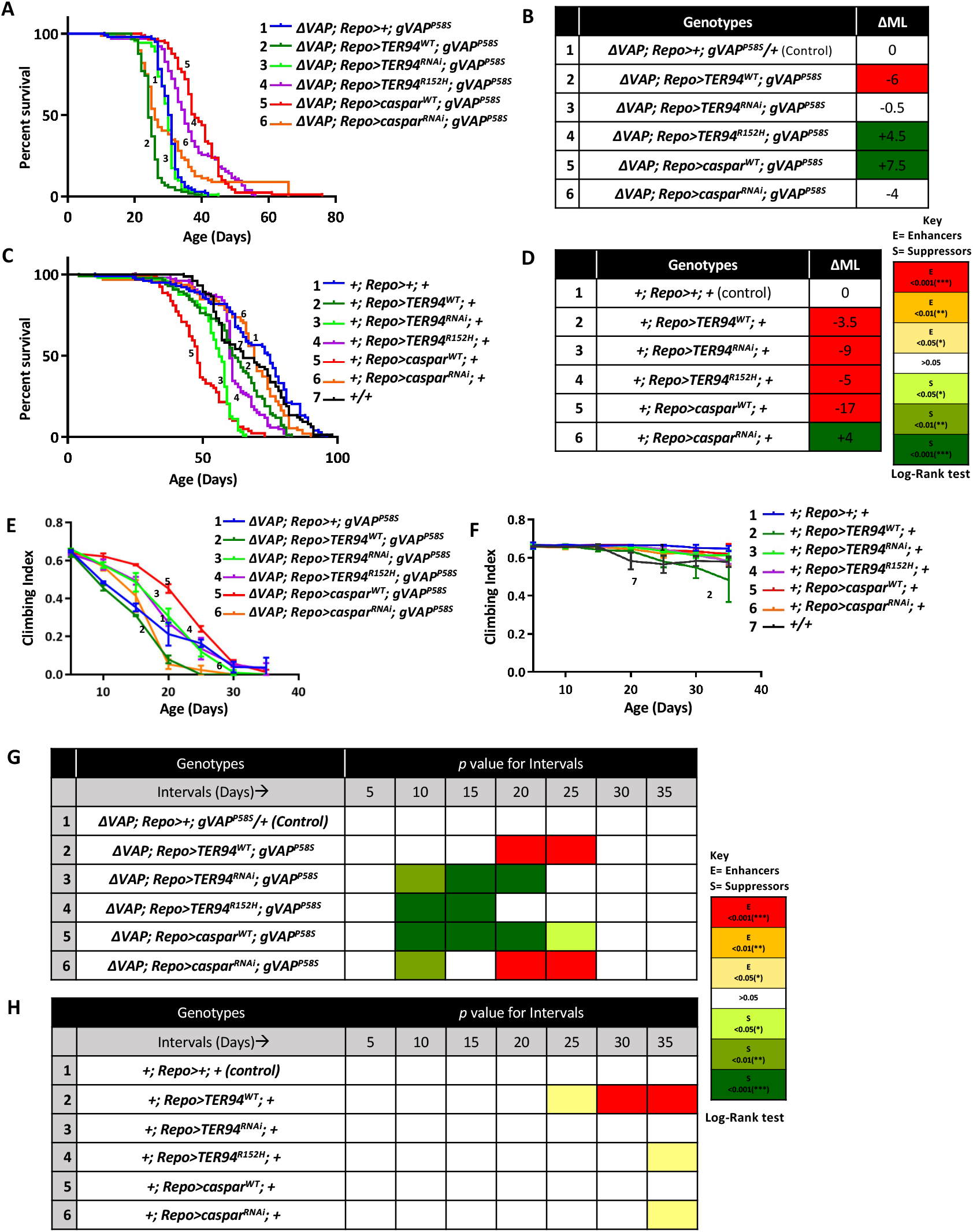
Overexpression of *caspar* in the glia increases lifespan and delays motor dysfunction in the disease model. **A**. Lifespan curves for overexpression of *TER94*^*WT*^ (2, Dark green curve), *TER94*^*RNAi*^ (3, light green), *TER94*^*R152H*^ (4, purple), *caspar*^*WT*^ (5, red) and *Caspar*^*RNAi*^ (6, orange). *∆VAP/Y; Repo>+; gVAP*^*P58S*^*/+* (1, blue, ML=30.5 days) was used as the control. n=∼100 flies for every genotype. Curve comparison was done using Log-rank test. Combined *p*-value for the whole set is <0.001. For both *TER94*^*R152H*^and *caspar*^*WT*^ overexpression, there is an increase in lifespan. **B**. Tabulated results for lifespan recordings in panel A, quantified in the form of ΔML. The colour codes are as described previously (Fig 1D). **C**. Lifespan curves for overexpression of *TER94*^*WT*^ (2, dark green curve), *TER94*^*RNAi*^ (3, light green), *TER94*^*R152H*^ (4, purple), *caspar*^*WT*^ (5, red) and *caspar*^*RNAi*^ (6, orange) using *+; Repo-Gal4; +* background. *+; Repo>+;+* (Curve 1 in blue color, ML= 75) was the control for curves 2-6, while *+/+* (wild type flies, black curve, 7) was used as the master control (ML= 65 Days) to compare with curve 1 (p>0.05, not significant). Figure C functions as the control for the respective gene specific expressions in Figure A. n=∼100 flies for every genotype. Curve comparison was done using the Log-rank test. Combined *p*-value for the whole set is <0.001. **D**. Tabulated results for lifespan recordings in panel C, quantified in the form of ΔML. The colour codes are as described previously. **E**. Climbing indices for glial OE of *TER94*^*WT*^ (2, dark green curve), *TER94*^*RNAi*^ (3, light green), *TER94*^*R152H*^ (4, purple), *caspar*^*WT*^ (5, red) and *caspar*^*RNAi*^ (6, orange). *∆VAP; Repo>+; gVAP*^*P58S*^*/+* (1, blue color) was used as the control. The individual *p*-values are listed in panel G. For both *TER94*^*R152H*^and *caspar*^*WT*^ OE, there is an age dependant improvement in motor function. **F**. Climbing indices for glial overexpression of *TER94*^*WT*^ (2, dark green curve), *TER94*^*RNAi*^ (3, light green), *TER94*^*R152H*^ (4, purple), *caspar*^*WT*^ (5, red) and *caspar*^*RNAi*^ (6, orange) in the wild type background. *+; Repo>+;+* (1, blue color) was the control for curve 2-6, The individual *p*-values are listed in panel H. +/+ (wild type flies, black curve, number 7) was used as the master control to compare curve 1, the *p*-values for this comparison were: Day 5, *p*>0.05 (ns, not significant), Day 10, *p*>0.05 (ns), Day 15, *p*>0.05 (ns), Day 20, *p*>0.01(**), Day 25, *p*>0.001(***), Day 30, *p*>0.05(*) and Day 35, *p*>0.05(*). **G**. Individual *p*-values for intervals (5-day) for the climbing index of the animals in the *∆VAP; Repo-Gal4; gVAP*^*P58S*^ background are plotted as a colour code. As compared to control (row 1), *caspar*^*WT*^ overexpression leads to improved motor activity in the age range 10-25 days, while for *TER94*^*R152H*^, the period is 10-15 days. **H**. Individual *p*-values for intervals (5-day) for the climbing index of the animals in the *Repo-Gal4* background. As compared to control (row 1), *caspar*^*WT*^ overexpression does not improve motor activity in the age range 1-30 days.

### Expression of Caspar in the glia delays age dependant motor deterioration

The *∆VAP; gVAP*^*P58S*^ fly line develops progressive motor defects (Moustaqim-Barrette *et al*. 2014)(Fig. 3E). Since *OE* of both the *TER94*^*R152H*^ and *caspar* increase the lifespan of the VAP^P58S^ disease model, we measured the changes seen in motor activity, using the ability of flies to climb as a readout. The data were measured and analysed as described (Materials & Methods) and displayed in in terms of a climbing index (Fig. 3E,F). *caspar* overexpression significantly improved motor function (Fig. 3E, 5-red line), when compared to the control (Fig. 3E, 1-dark blue curve). *TER94*^*R152H*^ *OE* also showed a mild improvement in motor function, at par with *TER94 KD*. In comparison, when expressed in a wild-type fly, motor function did not improve for any of the genotypes (Fig. 3F) tested, with the possible exception of TER94 *OE* (Fig. 3F, 2-Dark green curve). For a better appreciation of the age-dependence of motor function, we have plotted the data at five day intervals (Fig. 3G,H; *Suppl. Fig. 3A,B*). In Fig. 3 G,H the statistical significance is color coded, with yellow, orange and red signifying deterioration of motor function, and shades of green indicating improvement in motor function, for animals of equivalent age. For the disease model (Fig 3G), in the age group 10-20 days, there appears to be a significant improvement of motor function for *TER94*^*R152H*^, *TER94 KD* and *caspar OE*. In contrast, overexpression of *TER94*^*WT*^ and *caspar* RNAi led to a significant deterioration of motor function for flies aged 20 and 25 days. For wild type flies, *TER94*^*WT*^ *OE* was detrimental for motor function after 25 days (Fig. 3H).

Both the lifespan and motor dysfunction data suggest that the VAP:Caspar:TER94 complex in glia may be an important contributor for the progression of the disease. TER94 is an important member of the UPS, that leads to proteasomal degradation of proteins (Yeung *et al*. 2008; van den Boom and Meyer 2018). With Caspar having an ability to coordinate transfer of ubiquitinated proteins to TER94, and with VAP acting as a docking site for Caspar, there is a strong possibility that these proteins cooperate in a physiological function that is related to proteostasis. This common function may be the Endoplasmic-Reticulum Associated Degradation (ERAD)(Ruggiano *et al*. 2014; Frakes and Dillin 2017) or clearance of VAP^P58S^ inclusions/ aggresomes (Chaplot *et al*. 2019). To confirm the same, we measured the change in the inclusion status of VAP aggregates in the brain of the adult fly, in response to *caspar* overexpression, in an age dependant manner.

### Caspar expression does not modulate VAP^P58S^ inclusions in the adult brain

In an earlier study, working with the larval brain, we had uncovered the physiological basis of the genetic interaction between SOD1/ALS1 and VAP/ALS8 (Deivasigamani *et al*. 2014; Chaplot *et al*. 2019). Cellular Reactive Oxygen Species (ROS), when increased in response to SOD1 malfunction triggered proteasomal activity, which led to a clearance of VAP^P58S^ aggregates from brain cells. The density of VAP^P58S^ inclusions in the brain was measured and found to decrease significantly with increase in cellular ROS (Chaplot *et al*. 2019). A similar methodology was developed to measure aggregates in the adult fly brain of the disease model. Interestingly, the density or area of VAP inclusions in the brain did not increase or decrease significantly on *caspar* OE or even *caspar* KD in the glia (Fig 4), in both larval (*Suppl. Fig. 4*) and adult brains (Fig. 4). This would suggest that increased lifespan and improved motor function that we see are not related to the ‘average’ density or size of VAP^P58S^ inclusions, which can be visualized by microscopy. This suggests that the VAP:Caspar:TER94 function that we are attempting to elucidate may either modulate proteostasis of proteins other than VAP inclusions or it may suggest that the complex has other, unknown functions in the cell. Results similar to ours have been found in a mammalian cell culture model (Genevini *et al*. 2014).

**Figure 4.**
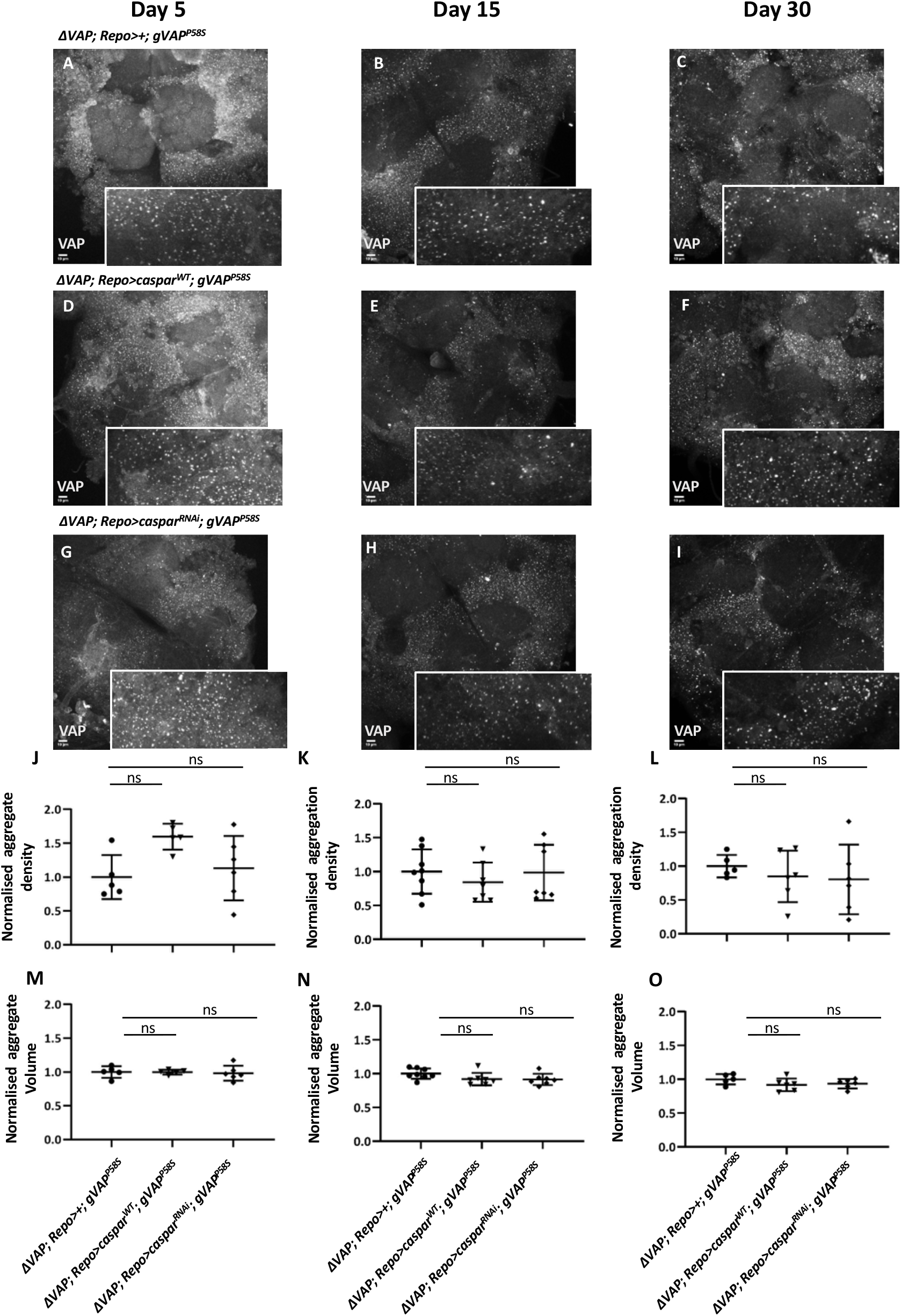
VAP inclusions in the brain of adult animals maintain *status-quo* in response to *caspar* overexpression. Representative images of VAP protein inclusions in adult brains for days 5, 15 and 30. The inclusions are marked with an anti-VAP antibody. The scale bar for the image is 10 μm. The inclusions are highlighted (2X digital zoom) in the inserted panel for each image. **A-C**. *∆VAP; Repo>+; gVAP*^*P58S*^. **D-F**. *∆VAP; Repo>caspar*^*RNAi*^; *gVAP*^*P58S*^ **G-I**. *∆VAP; Repo>caspar*^*WT*^; *gVAP*^*P58S*^ **J-O**. Graphical representation of normalised ‘aggregate density’ (J-L) and normalised ‘aggregate volume’ (M-O) of VAP inclusions, as defined in Materials and Methods. The genotypes compared are *∆VAP; Repo>+; gVAPP58S /+* (control), *∆VAP; Repo>casparWT; gVAP*^*P58S*^, and *∆VAP; Repo>caspar*^*RNAi*^; *gVAP*^*P58S*^. n= 5-10 brain samples. One-way ANOVA followed by Tukey’s multiple comparison (*P<0.05, ***P<0.001, ****P<0.0001; ns, not significant). Error bars indicate s.d.

The only known function in fly literature for Caspar is that it is a regulator of NFκB signalling (Kim *et al*. 2006). Caspar negatively regulates Immune Deficient (IMD)/NFκB signalling by controlling activity of the protease Death related ced-3/Nedd2-like caspase (Dredd), a Caspase-8 ortholog (Leulier *et al*. 2000; Stoven *et al*. 2000), which in turn cleaves the full-length precursor Relish (Rel), a fly NFκB, which is sequestered in the cytoplasm. FAF1 (Menges *et al*. 2009) has similar roles in NFκB signalling (Min-Young *et al*. 2004; Park *et al*. 2007), negatively regulating signalling by influencing the IκB kinase (IKK) complex. Overexpression of FAF1 in Jurkat cells causes cell death, a phenomenon that is dependent on its interaction with Fas associated death domain (FADD) and Caspase-8 (Ryu *et al*. 2003). Dredd was itself discovered as a novel effector for apoptosis (Chen *et al*. 1998) and is a FADD interactor (Hu and Yang 2000). Cleavage of Rel generates an N-terminal 68 kD polypeptide (REL68) that can be transported to the nucleus and transcriptionally activate defence genes. Increase in Caspar activity leads to reduction in IMD signalling while absence of Caspar leads to hyper-activation of IMD signalling and hence inflammation (Kim *et al*. 2006), in immune cells. A possible hypothesis for our disease model is that Caspar activity or levels may regulate import of REL68 in the glia, and that increased inflammation in the disease model, a consequence of reduced Caspar function may lead to a shorter lifespan and decreased motor function. If this hypothesis is true then two conditions need to be met. First, there should be increased inflammation in the disease model, especially for older flies, when compared to controls and second, the modulation of IMD/Rel signalling in glia should reduce or enhance the progression of the disease.

### Progression of ALS8 is modulated by the extent of glial inflammation

Healthy ageing in *Drosophila* includes the age-dependant upregulation of the immune response (Kounatidis *et al*. 2017), regulated by the IMD/Rel pathway and is strongest in the fly head (Kounatidis *et al*. 2017). We measured inflammation in the brain by measuring the levels of mRNA for Imd/Rel targets such as *attacinD, diptericin, drosocin* and *cecropinA1* and a Toll/Dif target *metchnikowin*, in the dissected heads of adult *Drosophila*, as a function of age. On day 5, *∆VAP; Repo>+; gVAP*^*P58S*^ heads had *diptericin and attacinD* levels with higher averages, but not statistically significant. For 5-day old animals, *drosocin* and *cecropinA1* levels were significantly higher than controls (Fig 5A). For 15-day old animals, where the disease had progressed, for *diptericin, drosocin* and *cecropinA1*, there is a 2-4-fold increase in mRNA levels as compared to controls. For *attacinD, diptericin* and *drosocin*, overexpression of *caspar* in the glia reduces levels of these antimicrobial peptide genes, by ∼2-fold. *metchnikowin*, a target of Toll signalling does not show age dependant inflammation and is not influenced by *caspar* overexpression (Fig 5A). Thus, on the 15^th^ day, with disease onset well underway, and with ∼80% of the animals still alive, there was a significant increase of inflammation in the head, which was reined in by Caspar.

**Figure 5.**
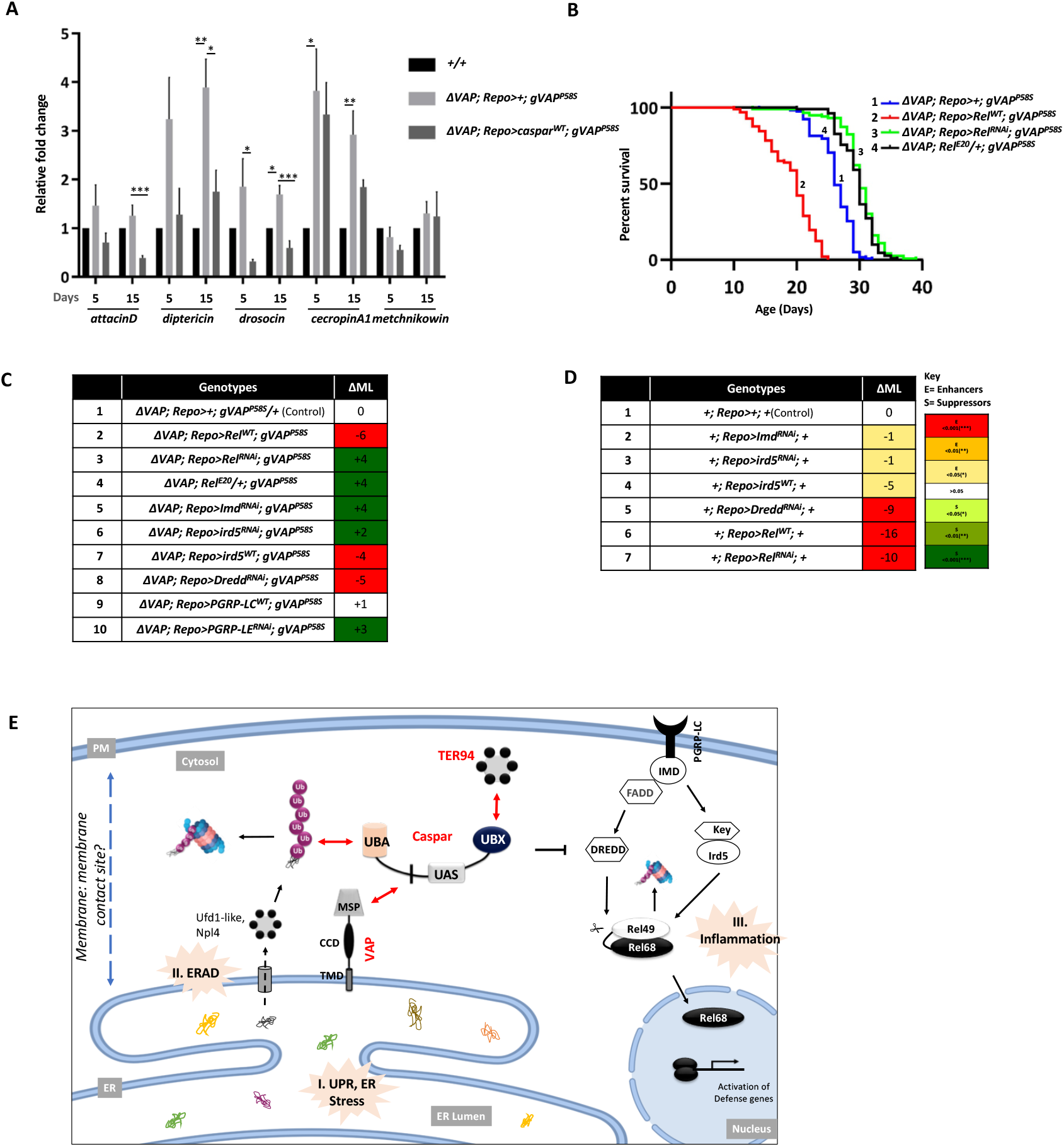
Inflammation in glia, regulated by IMD/Rel signalling contributes to the progression of the disease. **A**. Expression of Rel target genes *attacinD, diptericin, drosocin, cecropinA1* and Toll pathway target *metchnikowin* in day-5 and day-15 adult heads of wild type (*+/+*), *∆VAP; Repo>+; gVAP*^*P58S*^ and *∆VAP; Repo>caspar*^*WT*^; *gVAP*^*P58S*^ lines. Values on the Y-axis depict the fold change normalized to the house keeping gene *rp49*. Values shown are mean± SEM. N=3, n=3. Statistical analysis by two-way ANOVA followed by Tukeys multiple comparison test. *p< 0.05, **p< 0.01, ***p< 0.001. **B**. Lifespan plots for overexpression of *REL*^*WT*^ (2, red curve), *Rel*^*RNAi*^ (3, light green), *REL*^*E20*^ (4, black) in the *∆VAP; Repo-Gal4; gVAP*^*P58S*^ genetic background. *∆VAP; Repo>+; gVAP*^*P58S*^*/+* (1, blue curve) was used as the control. Curve comparison was done using log-rank test. Combined *p*-value for the whole set is <0.001. **C**. Tabulated results (∆ML) for lifespan recordings for experiments displayed in panel B and *Suppl. Fig 5A*. Levels of statistical confidence are color coded. ML for control, *∆VAP; Repo>+; gVAP*^*P58S*^*/+*, is 26 days. **D**. Tabulated results (∆ML) for lifespan recordings for controls as shown in *Suppl. Fig 5B*. Levels of statistical confidence are color coded. **E**. VAP^P58S^ disease model depicting the involvement of the VAP:Caspar:TER94 interaction in glial proteostasis, and its connectivity with IMD/Rel signalling, which regulates inflammation. The association of Caspar with both ER-based VAP and the PM based IMD/Rel Signalosome suggests that the ER:PM membrane contact sites may be the site for Caspar function. The key features of the model include (I) ER Stress caused directly or indirectly VAP misfolding (II) An up-regulation of the protein degradation machinery in response to ER Stress. VAP appears to act as a docking station, interacting with Caspar and TER94 to assist in proteasomal clearance. The AAA-ATPase TER94, itself an ALS locus, plays a central role for this function. (III) Age dependant Inflammation, caused by mis-regulation of Imd/Rel signalling. Our model suggests that Caspar’s function as a negative regulator for Imd/Rel signalling is attenuated in the disease. This may be a consequence of an increased partitioning of the available glial Caspar pool for VAP or TER94 related functions. As the adult animal ages, a tipping point is reached when the increasing burden of ER stress, proteasomal overload and glial inflammation leads to motor neuron cell death, followed by the death of the organism.

To test the second prediction, we modulated the levels of IMD/Rel signalling to upregulate or downregulate glial inflammation. Enhanced inflammation should lead to a reduction in ML and reduction of inflammation should increase the ML of the disease model. First, we measured the change in lifespan of *∆VAP; Repo>+; gVAP*^*P58S*^ flies after overexpressing and knocking down *Rel*, the transcriptional effector of the IMD/Rel pathway. *Rel OE* leads to increased inflammation and reduced lifespan by ∼6 days (Fig. 5B,C). *KD* of *Rel* by RNAi or reduction of *Rel* by ∼50% (*Rel*^*E20*^/+), increased ∆ML by ∼4 days (Fig. 5B,C). Next we tested other elements of the IMD/Rel pathway (Fig 5E). Knockdown of *IMD, IRD5* and *PGRP-LC* showed an enhancement in lifespan by 2-3 days (Fig. 5C), suggesting that these genes were part of a sterile inflammation cascade and/or a feedback loop. *OE* of IRD5 decreased lifespan (∆ML=-4), while *OE* of *PGRP-LC* did not modulate lifespan, whereas KD of *PGRP-LE* increased ∆ML by 3 days. Knockdown of *Dredd* reduced lifespan (∆ML=-5). The Dredd result was unexpected as a reduction in *Dredd* activity should reduce Rel signalling and reduce inflammation. Control lifespan experiments (*Suppl. Fig. 5*; Fig. 5D), in wild type flies, demonstrate that both *gain-of-function* and *loss-of-function* of *Rel* activity shortens lifespan as does reduction of *Dredd* in the glia. Dredd loss of function may have a role in triggering cell death and influence inflammation. The single available Dredd RNAi stock (BDSC, 34070) appears to be a weak and sick stock, and this may also confound the data. *Dredd* resides on the X chromosome, which makes it challenging to use *Dredd* deficiencies for our male specific readout. Another possibility is that Rel cleavage in glia may be Dredd independent, with an alternative pathway used for Rel cleavage and import. For example, in mammals, NFκB cleavage and maturation is a function of the proteasome (Sears *et al*. 1998), rather than a Dredd ortholog. Caspar, like FAF1 in mammals may directly bind to and influence Rel maturation.

In summary, IMD/Rel signalling (reviewed by (Lemaitre and Hoffmann 2007; Ligoxygakis 2013; Myllymaki *et al*. 2014; Zhai *et al*. 2018)), in the glia, appears to play a major role in regulating the lifespan of our disease model. Caspar can negatively regulate IMD/Rel signalling in glial cells and alleviates disease by reducing inflammation in the brain.

## DISCUSSION

Over the past decade it has become apparent that NFκB signal transduction cascades have diverse roles in the brain (reviewed by (Meffert and Baltimore 2005; Boersma and Meffert 2008; Kaltschmidt and Kaltschmidt 2009)). Major functions of NFκB signalling include the regulation of the neuronal immune response (Nguyen *et al*. 2002; Sochocka *et al*. 2017), learning and memory (Lubin and Sweatt 2007), neural development (Albensi and Mattson 2000; Beattie *et al*. 2002; Nickols *et al*. 2003; Rolls *et al*. 2007) and neuronal cell death (Chinchore *et al*. 2012). In flies, the Toll/ NFκB pathway has been linked to regulation of post-synaptic GluRIIA levels (Heckscher *et al*. 2007) while the IMD/ NFκB signalling appears to have important behavioural, apoptotic (Chinchore *et al*. 2012) and neuro-inflammatory roles (Cao *et al*. 2013; Petersen *et al*. 2013; Kounatidis *et al*. 2017; Li *et al*. 2018). In 2017, the Ligoxygakis lab (Kounatidis *et al*. 2017), found that IMD/NFκB signalling in the *Drosophila* brain was important for a normal lifespan, with negative regulators of IMD/NFκB playing critical roles in reducing inflammation in the brain. Enhanced inflammation was a key feature in age-dependant neurological decline. Reduction of inflammation, specifically in the glia could reverse phenotypes associated with age dependant neurodegeneration. In agreement with this idea, lifespan in a fly model of SCA (Li *et al*. 2018), where SCA3^polyQ78^ was expressed in neurons, could be extended by reduction of IMD/Rel signalling. Earlier, in a SOD1^G93A^ mouse model for ALS, it was found that NFκB signalling is upregulated in spinal cords, a feature also seen in the same tissue in human patients (Frakes *et al*. 2014). In the mouse model, cessation of NFκB activity (Frakes *et al*. 2014) in microglia rescued motor neurons from early death and extended lifespan by reduction of inflammation. Glia have thus emerged as a cell type where NFκB has critical roles in both vertebrates and invertebrates (Bazan 2009; Kounatidis and Chtarbanova 2018). C9orf72, a gene that is found to be perturbed in 60% of sALS patients appears to be required for proper macrophage and microglial function in mice (O’Rourke *et al*. 2016). Also, NFκB signalling appears to be activated in both fALS and sALS (Swarup *et al*. 2011). NF-κB, thus appears to be an important signalling pathway in the glia (ONeill and Kaltschmidt 1997; Beattie *et al*. 2002).

Our understanding of the mechanisms that fine tune IMD/Rel signalling has grown in leaps and bounds in the last decade (Lemaitre and Hoffmann 2007; Myllymaki *et al*. 2014; Kleino and Silverman 2019). An emerging theme in NFκB signal transduction is the detection of large cytoplasmic supramolecular protein complexes that undergo clustering with a evidence of amyloid formation and colloidal phase separation (Dickens *et al*. 2012; Fu *et al*. 2016; Kleino *et al*. 2017; Kleino and Silverman 2019). These compartments increase the local concentration of the elements involved in signalling, and their formation and stability in response to immune signalling depends on dynamic post translational modifications (PTMs), with poly-ubiquitin chains playing a central role (Zhou *et al*. 2005; Paquette *et al*. 2010; Meinander *et al*. 2012; Chen *et al*. 2017), possibly offering docking sites for complex formation. A feature of the NFκB ‘signalosome’ is the clustering of IMD, FADD and Dredd at the PGRP-LC receptor, near the plasma membrane (PM). In *Drosophila*, the death effector domains and the death interaction domains are binding partners for the formation of one of the core components of the complex (Hu and Yang 2000; Kleino and Silverman 2019). The association of IMD/Dredd/Caspar/FADD via these domains has critical roles in modulating signalling for both host defence and cell death (Georgel *et al*. 2001; Chinchore *et al*. 2012; Kim *et al*. 2014).

Our study puts the spotlight on Caspar and its role in the cell, potentially as adapter protein that connects plasma membrane (PM)-based PGRP-LC amyloid clusters to the VAP domain at the ER. An extension of the above idea would suggest that Caspar may perform its function in membrane contact sites (MCS), where the PM and ER are in close association (Zaman *et al*. 2020). VAP, itself is enriched in MCS, interacting with partner proteins in opposing membranes to maintain membrane: membrane spacing (Murphy and Levine 2016; Yadav *et al*. 2018). The spatially restricted MCS, along with the clustering of IMD/Rel pathway proteins would form an ideal compartment for regulated signalling. Intriguingly, VAP has earlier been identified as a physical interactor of Rel (Fukuyama *et al*. 2013). This study from the Hoffmann lab uncovered 369 protein:protein interactions in the IMD/Rel pathway, again underscoring the role of protein:protein interactions in IMD/Rel signaling. In the same study, Ird5 was found to be a conjugation target for the PTM, SUMO. The SUMO conjugation resistant variant Ird5^K152A^ was found not to be an efficient transducer of the immune signal. Caspar itself is SUMO conjugated at K551 (Handu *et al*. 2015). In addition to ubiquitination and phosphorylation, SUMO conjugation may play an important role (Hegde *et al*. 2020) in formation of signalosomes, especially with many of the proteins involved containing SUMO interaction motifs. The PM:ER MCS could serve as a domain where the VAP:Caspar:TER94 complex could associate with proteins that are part of the IMD/Rel signal transduction cascade. Caspar, with its ability to bind to poly-Ub proteins, to TER94 and to VAP has the potential to bring together VAP with functional elements of the proteasome and the NFκB signalosome.

An important question that remains is the molecular function of Caspar in the cell. Based on the available data there are two possibilities. One, Caspar may be a facilitator of protein degradation and in addition to assisting in VAP mediated protein degradation, it may regulate IMD/Rel signalling by actively degrading proteins, which would include targeting elements such as Dredd and Rel for degradation. The exact mechanism for regulation of Dredd by Caspar (Kim *et al*. 2006) has not been worked out and its degradative function may well be the answer. Two, Caspar may be an adapter that regulates the association of signalling complexes. For the second function, it may assist in capturing poly-Ub chains for modification of proteins in the signalosome. Activation via Ub-conjugation is a theme in activation of the pathway (Paquette *et al*. 2010; Meinander *et al*. 2012). The Silverman and Meier labs have earlier documented mechanisms involved for K63-polyUb conjugation of IMD and Dredd by the E3-ligase DIAP2, along with the accessory proteins Effete, Bend and Uv1a, all of which are part of the complex (Paquette *et al*. 2010; Meinander *et al*. 2012). Active control of IMD/Rel signalling may require ER:PM contact with many of the components discussed above involved in fine-tuning the signal.

Our model (Fig. 5E), which should parallel the onset and progression of disease in ALS incorporates the three major physiological events that are central to glial homeostasis. First, animals expressing VAP^P58S^, in the absence of wild-type VAP show ER stress (Kanekura *et al*. 2009; Mori *et al*. 2011; Moustaqim-Barrette *et al*. 2014; Larroquette *et al*. 2015) and morphological changes in the ER (Fasana *et al*. 2010; Kuijpers *et al*. 2013), a direct or indirect response to misfolded or partially-folded mutant VAP proteins. The fact that *VAP*^*P58S*^ lines can progress through development, in the absence of *VAP*^*WT*^, suggests that VAP^P58S^ is partially active, but this allelic variant is unable to sustain adult life beyond ∼25 days. Second, the ER stress should lead to enhanced clearance by the UPS, with pathways such as ERAD (Vembar and Brodsky 2008), working overtime to maintain proteostasis. The evidence for VAP and Caspar’s involvement in proteasomal clearance is primarily based on its interaction with TER94 and the presence of ERAD components Ufd1-like and Npl4 in Caspar IPs. This in turn could lead to overload at the proteasome (Moumen *et al*. 2011), though experiments with CDC3δ suggest otherwise (Genevini *et al*. 2014). Last, but not the least, there appears to be an age dependant increase in glial inflammation, which can be reined in by an increase in Caspar levels. This suggests that the pool of Caspar available to negatively regulate Rel is reduced in our ALS disease model. Logically, the reason for this decrease would be the sequestration of Caspar, either by VAP^P58S^ or by the strained proteasomal machinery.

In summary, we find that Caspar regulates disease progression, primarily by regulating inflammation in glia. VAP appears to function as a docking site for Caspar; Caspar connects not only two ALS orthologous loci, namely VAP and TER94, together, it also links these to IMD/Rel signalling. Caspar is therefore a point of convergence for critical cellular pathways that are central to motor neuron disease.

## MATERIALS & METHODS

### *Drosophila* husbandry, Stocks and reagents

All flies were raised and crosses were conducted at 25°C in standard corn meal agar. The flies expressing *genomic* VAP^WT^ (*gVAP*^*P58S*^ *(VK31)/TM3TB*) and VAP^P58S^ (*gVAP*^*WT*^ *(VK31)/TM3TB*) were a kind gift from Hiroshi Tsuda (Moustaqim-Barrette et al., 2013). We balanced these with *VAP∆166*, to generate *∆166/FM7A;+; gVAP*^*P58S*^ *(VK31)/TM3TB and ∆166/FM7A;+; gVAP*^*WT*^ *(VK31)/TM3TB*. The *VAP∆166* allele was used instead of *VAP∆31* used earlier because we were unsuccessful in receiving live *VAP∆31* flies from the Tsuda lab. *VAPΔ166 (P**ennetta* *et al. 2002)*, like the *VAPΔ31 allele*, is a larval/pupal lethal. The primary lines used for our experiments, *Δ166;+;gVAP*^*WT*^ and *Δ166;+;gVAP*^*P58*S^ are at par with the Tsuda lines in terms of lifespan, cytoplasmic VAP inclusions and motor dysfunction.

Further, these lines were modified by adding Gal4 driver chromosomes for the second chromosome. The Gal4 drivers used were *MHC-Gal4, OK6-Gal4* and *Repo-Gal4*. The *UAS-VAP*^*WT*^ and *UAS-VAP*^*P58S*^ were generated in the Jackson lab (Ratnaparkhi *et al*. 2008). Lines procured from the Bloomington *Drosophila* Stock Centre are: Name (Stock #), Canton-S (0001), *UAS-VAP*^*RNAi*^ *(27312), UAS-TBPH*^*RNAi*^(29517), *UAS-caz*^*WT*^ (17010), *UAS-caz*^*RNAi*^ (34839), *UAS-SOD*^*WT*^(24754), *UAS-SOD1*^*RNAi*^ (34616), *UAS-TER94*^*RNAi*^ (32869), *UAS-alsin*^*WT*^ (27162), *UAS-alsin*^*RNAi*^ (28533), *UAS-senataxin*^*RNAi*^ (34683), *MHC-Gal4* (38464), *OK-Gal4* (64199), (*UAS-caspar*^*RNAi*^(44027), *UAS-Rel*^*RNAi*^ (33661), *UAS-Rel*^*WT*^ (9459), *Relish*^*E20*^ (55714), *UAS-Imd*^*RNAi*^ (38933), *UAS-ird5*^*WT*^ (90312), *UAS-ird5*^*RNAi*^ (57751), *UAS-Dredd*^*RNAi*^ (34070), *UAS-PGRP-LE*^*RNAi*^ (60038), *UAS-PGRP-LC*^*WT*^(30919), *UAS-caspar*^*RNAi*^(44027), *caspar*^*lof*^ (11373), *caspar* deficiency (23691. Lines procured from the National Institute of Genetics were *UAS-Rel*^*11992R-1*^ and *UAS-Rel*^*11992R-2*^. The *UAS-TBPH:FLAG:HA*^*WT*^ line was procured from the National Centre for Biological Sciences Stock Centre (NCBS,(Diaper *et al*. 2013)), while *UAS-TER94*^*WT*^, *UAS-TER94*^*R152H*^, *UAS-TER94*^*A229E*^ were procured from the Taylor laboratory (Ritson *et al*. 2010). Third chromosome *apterous-Gal4* and *daughterless* Gal4 were procured from the Shashidhara laboratory. The second chromosome *Repo-Gal4* (Lee and Jones 2005), was a kind gift from Dr. Bradley Jones. *UASt-caspar:HA* was generated by procuring the cloned vector UFO05904 from the *Drosophila* Genomics Resource Centre tagged-ORF collection and generating transgenic animals at the NCBS transgenic facility. Expression of the line was validated by monitoring transcripts by real-time PCR and protein expression by anti-HA and anti-Caspar antibodies. The *caspar*^*lof*^ (*casp*^*c04227*^) line is a gypsy insert in the *caspar* locus and is a strong hypomorphic (near-null) allele (*Suppl. Fig. 2A*).

### Lifespan and survival analysis

Survival assays were carried out on the *genomic* VAP (*gVAP*) lines (Fig 1A) and their derivatives (*Suppl. Fig 1*; Fig. 1D). For each experiment, ∼100 F1 male flies of the desired genotype were collected with each vial containing 15 or less age-matched flies. Animals were flipped to a fresh vial every fourth day, with number of flies recorded per vial on a daily basis, till all flies, including those in control experiments were dead. The survival data was plotted and analysed using the log-rank test in Prism 7, which compares the entire survival curve(s) and a gives a value of significance as a p-value, which was recorded and reported as a colour-coded map (Fig. 1D). The ML for each curve was also calculated and used as a simple readout for each experiment.

### Generation of Caspar Antibody

Full length *caspar* was sub-cloned into pET-45b. The N-terminal 6X-His-Tagged Caspar was expressed in *E. coli* BL21DE3 cells. 1mM Isopropyl β-D-1-thiogalactopyranoside was used to induce expression of protein at 25 °C. The cellular lysate, in 1X TBS, 10 mM Imadizole was incubated with Ni-NTA beads (Qiagen) and the bound protein of interest was eluted using increasing concentrations of Imidazole, namely, 25 mM, 50 mM, 100 mM, 150 mM. Caspar eluted at both 50 and 100 mM. The fractions were merged, concentrated and injected into a GE Healthcare Sephadex-G200 preparative column. Fractions of the major peak at 150 kDa were collected and eluted protein showed a molecular weight of 66 kDa in SDS-PAGE gels, suggesting that Caspar was a dimer in its native state. The column-purified protein was used to generate a Rabbit Polyclonal antibody by Bioklone (Chennai, India). The serum was further purified with Caspar conjugated to Protein A beads and the Caspar enriched IgG used for further experiments. The antibody was validated (*Suppl. Fig. 2*). The working dilutions for the antibody were 1:20,000 for Western Blots and 1:1000 for tissue.

### Immunoprecipitation, Western Blot Analysis

5-10 day old adult flies were lysed in Co-IP Lysis Buffer (20mM Tris pH 8.0, 137mM NaCl, 1% IGEPAL, 2mM EDTA, 1X PIC) using a Dounce homogenizer and centrifuged at 21,000*g* for 30 minutes. 3 mg of total lysate was incubated with 5μg of primary antibody (Rb anti-Caspar or Rb anti-VAP) and 5μg of Normal Rabbit IgG overnight at 4 °C. The Anti-VAP antibody was generated as described (Yadav *et al*. 2018; Chaplot *et al*. 2019). Antigen-antibody complexes were captured using 50μl of BioRad SureBeads Protein A (1614013) at 4 °C for 4 hours. Beads were washed six times with Co-IP Lysis Buffer and protein complexes eluted by boiling in 1X Laemmli Sample Buffer. Eluted proteins were resolved on a 10% polyacrylamide gel followed by western blotting or in-gel trypsin digestion. Proteins separated by SDS-PAGE were transferred onto a PVDF membrane (Immobilon-E, Merck) and blocked in 5% milk in Tris-Buffer Saline (TBS) with 0.1% Tween 20 (TBS-T) for an hour. Blots were then incubated overnight with primary antibody diluted in 5% milk in TBS-T, at 4 °C. Following three washes with TBS-T, blots were incubated with secondary antibodies diluted in 5% milk in TBS-T, for 1 hour at RT. Blots were washed thrice with TBS-T and visualized on a LAS4000 Fuji imaging system after incubating with Immobilon Western Chemiluminescent HRP substrate (Merck). The following antibodies were used: Rabbit anti-VAP, 1:10000 (Yadav *et al*. 2018; Chaplot *et al*. 2019); Mouse anti-α-Tubulin, 1:10000 (T6074, Sigma-Aldrich); Goat anti-rabbit HRP and Goat anti-mouse HRP secondary antibodies, each at 1:10000 (Jackson ImmunoResearch).

### In-gel Trypsin Digestion and LC-MS/MS Analysis

Before in-gel trypsin digestion of the Co-IP eluate, the antibody was crosslinked to the SureBeads using DMP (Sigma) according to the NEB crosslinking protocol to avoid elution of the antibody. After crosslinking 10μg Caspar antibody, Co-IP was performed as described above. In-gel trypsin digestion was carried out as previously described (Shevchenko *et al*. 2006). Briefly, Coomassie-stained bands on the gel were excised and cut into 1 mm cubes. Gel pieces were transferred to a clean microcentrifuge tube and destained with buffer containing 50% acetonitrile in 50 mM Ammonium bicarbonate. Reduction and alkylation were carried out on the destained gel pieces by incubating with 10mM dithiothreitol (DTT) followed by incubating with 20mM iodoacetamide. Gel pieces were saturated with sequencing grade Trypsin (Promega) at a concentration of 10 ng/μl and incubated overnight at 37 °C. Peptides were extracted by sequential addition of 100 μl of 0.4% Trifluoroacetic acid (TFA) in 10% ACN, 100 μl of 0.4% TFA in 60% ACN and 100 μl of ACN. The pooled extract was dried in a vacuum centrifuge and reconstituted with 50 μl of 0.1% TFA. The peptides in TFA were purified using the StageTip protocol (Rappsilber *et al*. 2007).

LC–MS/MS analysis was performed on the Sciex TripleTOF6600 mass spectrometer interfaced with an Eksigent nano-LC 425. Tryptic peptides (1 μg) were loaded onto an Eksigent C18 trap (5 μg capacity) and subsequently eluted with a linear acetonitrile gradient on an Eksigent C18 analytical column (15 cm × 75-μm internal diameter). A typical LC run lasted 2 h post loading onto the trap at a constant flow rate of 300 nL/min with solvent A consisting of water + 0.1% formic acid and solvent B consisting of acetonitrile. The gradient schedule for the LC run was 5% (vol/vol) B for 10 min, a linear gradient of B from 0% to 80% (vol/vol) over 80 min, 80% (vol/vol) B for 15 min and equilibration with 5% (vol/vol) B for 15 min. Data was acquired in an information-dependent acquisition (IDA) mode over a mass range of 300–2,000 *m*/*z*. Each full MS survey scan was followed by MS/MS of the 15 most intense peptides. Dynamic exclusion was enabled for all experiments (repeat count 1; exclusion duration 6 s). Peptide identification and quantification were carried out with the SCIEX ProteinPilot software at a false discovery rate (FDR) of 1%. A RefSeq *Drosophila* protein database (release 6) was used for peptide identification. Proteins that were identified in two or more replicates and had two or more quantified peptides were tabulated.

### Motor Function

Motor performance of each genotype was analyzed using the standard startle induced negative geotaxis climbing assay (Branco *et al*. 2008; Madabattula *et al*. 2015), with minor modifications. Three separate sets (biological replicates) of 30 age-matched adult males of each genotype were raised at 25 °C, with 15 males in each vial. Each experimental set of 30 flies was transferred in a 250 ml (30 cm) glass cylinder and allowed to acclimatise for around 5 minutes. These flies were then tapped to induce a startle and were allowed to climb for 60 s. After 60 s, the flies were scored into three kinds based on their position in the cylinder. Flies which did not climb were scored 0 (Non-Climbers), flies which climbed till 80 mL (7.5 cm) mark were scored 1 (Bad Climbers) and flies which climbed beyond 80 mL were scored 2 (Good Climbers). This was repeated thrice for each set of flies. A three minute resting window was used in between the trials for each genotype. The climbing assays were conducted for a genotype every 5 days till at least one genotype completely stopped climbing or the flies were dead. The flies were exposed to CO2 only after a day’s trial was complete, ensuring no effect of anaesthesia on the assay. Flies were transferred to a fresh vial every four days to avoid death due to sticky media. The scores were used to calculate the climbing index of replicates. Statistical analysis was performed using Two-way ANOVA followed by multiple comparison testing by Tukey test. The Climbing index is a proxy for the fitness of a particular fly/genotype on a particular day and can be used as a readout for any progression of the motor defect in a set of flies. The climbing index was calculated as described by (Azuma *et al*. 2014).

### Microscopy, Staining & Image analysis

For larval brains, wandering third instar larvae were selected and the brains dissected in phosphate buffer Saline (1X PBS). Fixation was carried out in 4%PFA with 0.3%Triton-X in PBS for 20 minutes followed by three washes with 1X PBS. The brains were blocked in 2% BSA with 0.3%Triton X-100 in PBS for 1.5-hours and then incubated in primary antibodies overnight at 4°C. This was followed by 1.5-hour wash with blocking solution and incubation in Alexa Fluor secondary antibodies for 1.5 hours at room temperature, followed by four 20 minute washes with the blocking solution. DAPI (4,6 diamidino-2-phenylindole DAPI) was added during the second wash (1:1000). Samples were then washed with PBS, mounted in 70% glycerol with n-propyl gallate and the areas of the ventral nerve cord were imaged under Zeiss Confocal microscopes at 63X oil magnification. Antibodies utilized are as follows: M Anti-Repo (DSHB; 1:100), Rat Anti-ELAV (DSHB;1:100), Rb Anti-VAP (Yadav *et al*. 2018; Chaplot *et al*. 2019). The analysis was as described earlier (Chaplot *et al*. 2019).

For the adult brains preparations, flies were anaesthetized using CO2 and their brains dissected in 1X PBS (pH 7.4). The brains were fixed in 1.2% PFA for 24 hours at 4°C, then washed twice in 5% PBST for 20 minutes, followed by two 30-minute washes in PAT (0.5% BSA with 0.5% Triton X-100 in 1X PBS) buffer. The samples were then blocked in 5% BSA for two hours. The brains were incubated in primary antibody for 36 hours at 4 °C then washed in PAT Buffer, followed by incubation with secondary antibody for another 36 hours. The samples were washed four times in PAT buffer, 1 hour each wash. DAPI was added during the second wash. Next, samples were washed with PBS, mounted in Slowfade mounting medium (Vectashield, S36937) and imaged. Images were acquired on a Leica Sp8, using a 63x objective with a zoom factor of 0.75. Z stacks were acquired with an interval of 1 micron at 16-bit depth. Images were analyzed by Huygens Professional software (HuPro CE 18.10.0p2) from Scientific Volume Imaging, as described (Chaplot *et al*. 2019), with modifications. 5-10 brains were analyzed per experiment with five regions of interest (ROI) per brain. The ROIs were chosen in the sub-esophageal region of the brain. Parameters measured were ‘aggregate volume’ in μm^3^ and ‘aggregate density’ as number of aggregates per μm^3^. Data was represented as normalized values to the control (*ΔVAP;Repo>+; gVAP*^*P58S*^) on each day (Fig. 4 J-O). Graphpad prism 7 was used to plot the data. For statistical analysis, one-way ANOVA was used followed by Tukey’s multiple comparison test statistical significance.

### Real-Time PCR

mRNA was extracted from 50-80 adult heads (5-day and 15-day old) using the Qiagen RNAeasy mini kit (74104). 1ug of mRNA was used for the cDNA synthesis using the High Capacity cDNA Reverse Transcriptase Kit (4368814) by Applied Biosystems. The qPCR reaction was carried out using KAPA SYBR FAST (KK4602) by Sigma using Eppendorf Realplex Mastercycler. The experiments were carried out thrice with three technical replicates each. The relative fold change for each genotype was calculated by normalizing to the housekeeping gene *rp49*. The data was the analysed using Two-way ANOVA followed by Tukey’s test for multiple comparison. Primers pairs used *(H**andu* *et al. 2015; K**ounatidis* *et al. 2017)* were as follows: *rp49 Forward(-f): GACGCTTCAAGGGACAGTATC, rp49 reverse(-r): AAACGCGGTTCTGCATGAG; attacinD-f: CGGTCAACGCCAATGGTCAT, attacinD-r: CATTCAGAGCGGCGTTATTG; diptericin-f: ACCGCAGTACCCACTCAATC, diptericin-r: GGTCCACACCTTCTGGTGAC; cecropinA1-f: CATTGGACAATCGGAAGCTGGGTG, cecropinA1-r: TAATCATCGTGGTCAACCTCGGGC; drosocin-f: GTTCACCATCGTTTTCC, drosocin-r: CCACACCCATGGCAAAAAC; metchnikowin-f: GCTACATCAGTGCTGGCAGA, metchnikowin-r: AATAAATTGGACCCGGTCT*.

## ACKNOWLEDGEMENTS

<u>We thank</u>: BDSC, supported by NIH grant P40OD018537, for fly stocks; FlyBase, supported by a grant from the National Human Genome Research Institute at the U.S. National Institutes of Health U41HG000739; National Institute Genetics (NIG), Japan, for fly stocks; Drosophila Genome Resource Centre (DGRC), supported by NIH grant 2P40OD010949, for vectors and clones; Transgenic RNAi project (TRiP), at the Harvard Medical School supported by NIH/NIGMS R01-GM084947 for providing transgenic RNAi fly stocks; Transgenic injection facility at the National Centre for Biological Sciences (NCBS); IISER Imaging Facility for access to microscopy and analysis resources; IISER Proteomics facility for training of students and access to instruments; Neel Wagh for technical support.

## Author Contributions

ST and GR conceptualized the project. ST, SH, AT, LG, BK executed the experiments, collected data and analysed the same. AR and GR were involved in supervision, project administration and funding acquisition. All authors contributed to the writing and editing of manuscript.

