## Supplementary files for "Caspar, an adapter for VAP and TER94 delays progression of disease by regulating glial inflammation in a *Drosophila* model of ALS8"

**Suppl. Figure 1. A lifespan based screen for genetic interaction between ALS orthologous loci in neurons, muscle and glia.** Tissue specific expression in muscle (*ΔVAP; MHC-Gal4; gVAP*$^{P58S}$), glia (*ΔVAP; Repo-Gal4; gVAP*$^{P58S}$*)* and motor neurons (*ΔVAP; OK6-Gal4; gVAP*$^{P58S}$) was used to overexpress and knockdown *VAP, TBPH, caz, SOD1, TER94*, *alsin* and *senataxin* in a *VAP*$^{P58S}$ genetic background.
Survival plots for muscle (*A-L*, Panel 1), Glia (*A-L*; Panel 2) and Neurons (*A-L*; Panel 3) are displayed for *VAP, TBPH, caz, SOD1, TER94, alsin* and *senataxin*. The blue curve represents the *ΔVAP; Gal4>+; gVAP*$^{P58S}$ which is the control and is constant in every graph. The red curve represents the locus studied and differs in every graph. The p-value from the log-rank test mentioned in each graph is the comparison of the red curve with the blue curve in each graph.

<u>Muscle screen</u>: The median value for *VAP*$^{WT}$= 21, *VAP*$^{RNAi}$= 23, *TBPH*$^{WT}$= 21, *TBPH*$^{RNAi}$= 23, *caz*$^{RNAi}$= 23, *SOD1*$^{WT}$= 21, *SOD1*$^{RNAi}$= 21, *TER94*$^{WT}$= 21, *TER94*$^{RNAi}$= 20, *alsin*$^{WT}$= 24, *alsin*$^{RNAi}$= 23, *senataxin* $^{RNAi}$= 22. Control: *ΔVAP; MHC>+; gVAP*$^{P58S}$ has a median life span of 26 days.
<u>Glial Screen</u>: The median value for *VAP*$^{WT}$= 23, *VAP*$^{RNAi}$= 26, *TBPH*$^{WT}$= 27, *TBPH*$^{RNAi}$= 27, *caz*$^{RNAi}$= 31, *SOD1*$^{WT}$= 25, *SOD1*$^{RNAi}$= 29, *TER94*$^{WT}$= 21, *TER94*$^{RNAi}$= 30, *alsin*$^{WT}$= 25, *alsin*$^{RNAi}$= 30, *senataxin*$^{RNAi}$= 30. Control: *ΔVAP; Repo>+; gVAP*$^{P58S}$ has a median life span of 30 days.
<u>Motor Neuronal Screen</u>: The median value for *VAP*$^{WT}$= 17.5, *VAP*$^{RNAi}$= 20 *TBPH*$^{WT}$= 18, *TBPH*$^{RNAi}$= 19, *caz*$^{RNAi}$= 20, *SOD1*$^{WT}$= 19, *SOD1*$^{RNAi}$= 17, *TER94*$^{WT}$= 17, *TER94*$^{RNAi}$= 17, *alsin*$^{WT}$= 19, *alsin*$^{RNAi}$= 18, *senataxin*$^{RNAi}$= 17. Control: *ΔVAP; OK6>+; gVAP*$^{P58S}$ has a median life span of 16 days.

The *caz* overexpression construct is an insert on the X chromosome of the fly line and hence could not be used for our male specific assay. 'OE' stands for overexpression and 'KD' stands for knockdown via RNA interference.

Suppl. FIGURE 1

#### *MHC-Gal4* (Panel 1)

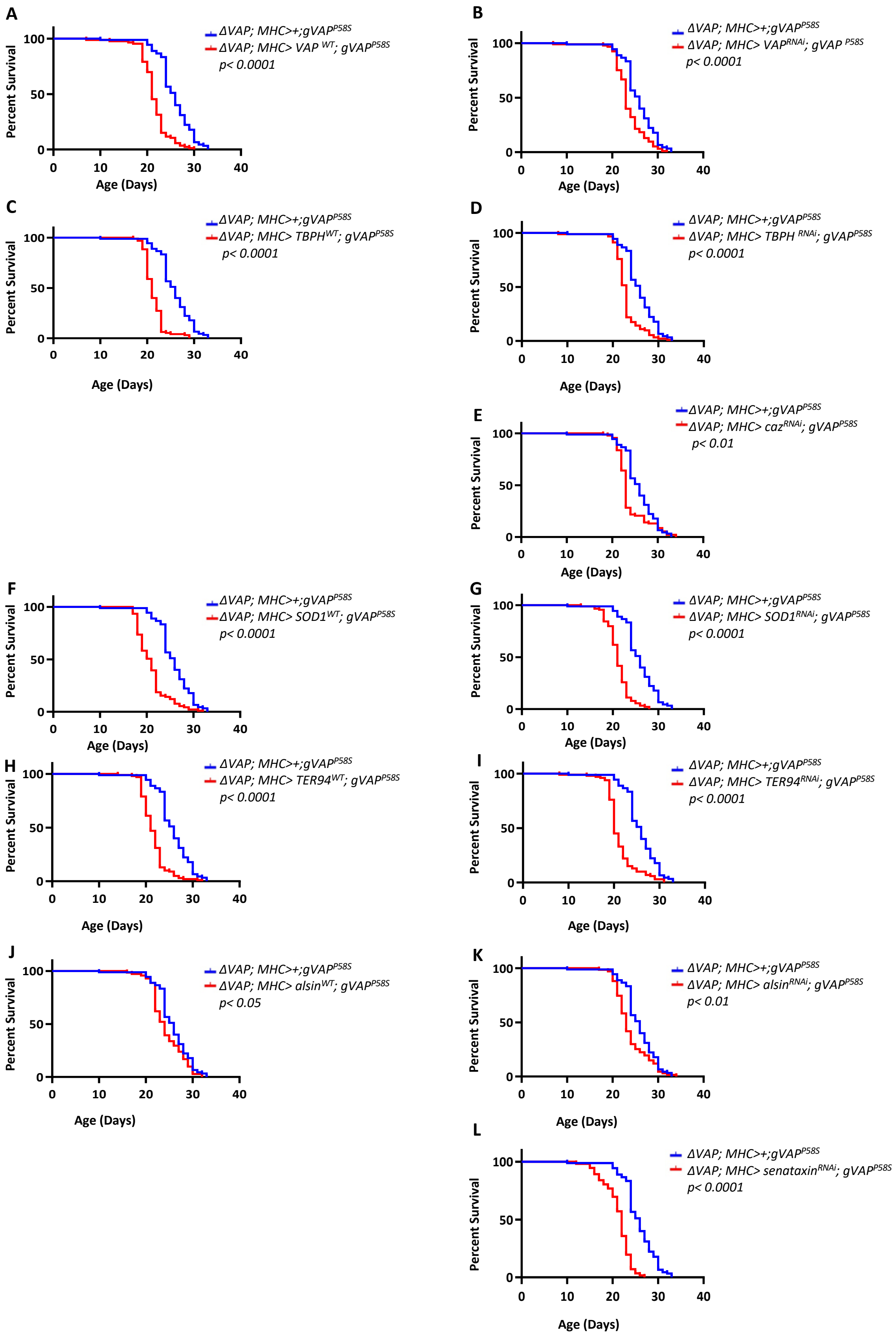

Suppl. FIGURE 1

### *Repo-Gal4* (Panel 2)

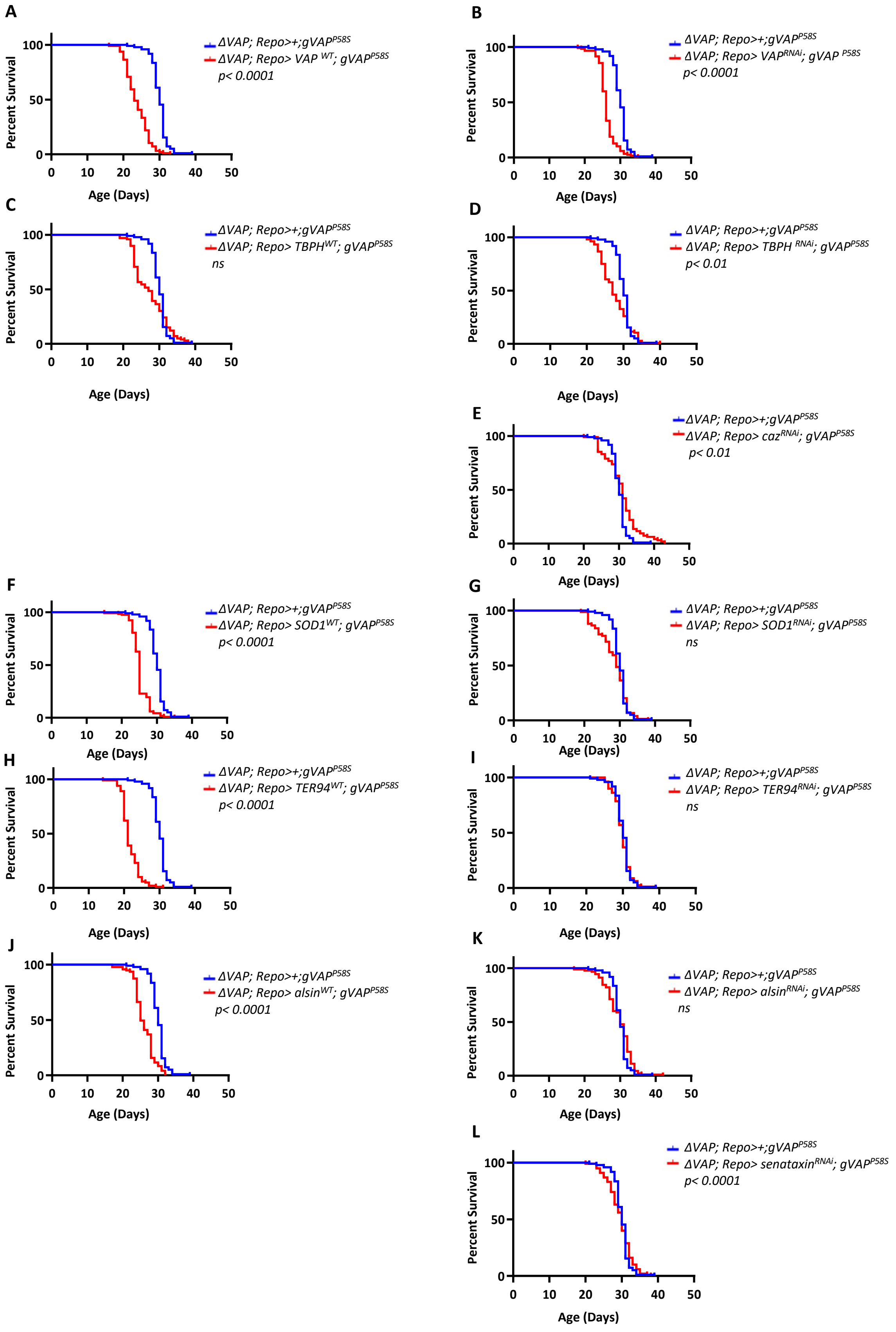

### *OK6-Gal4* (Panel 3)

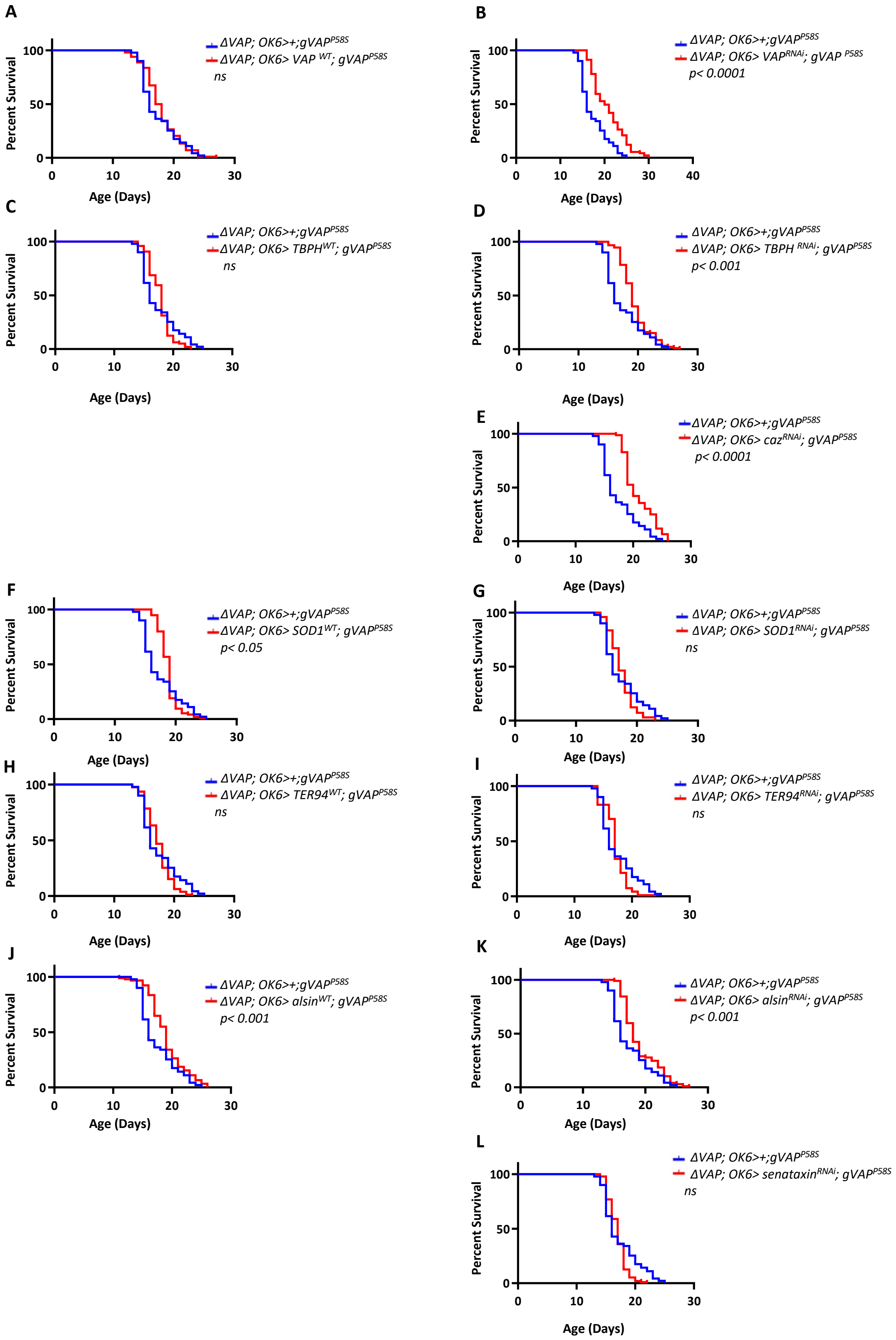

Suppl. FIGURE 2

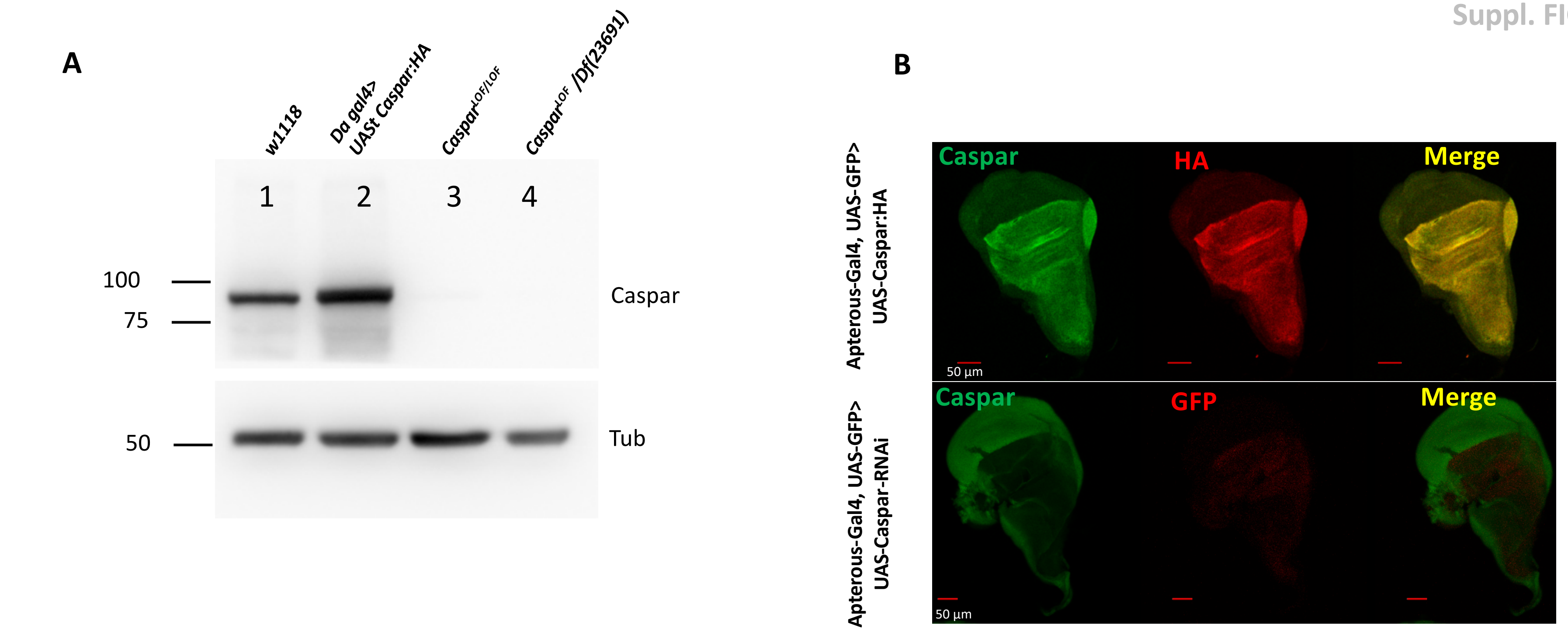


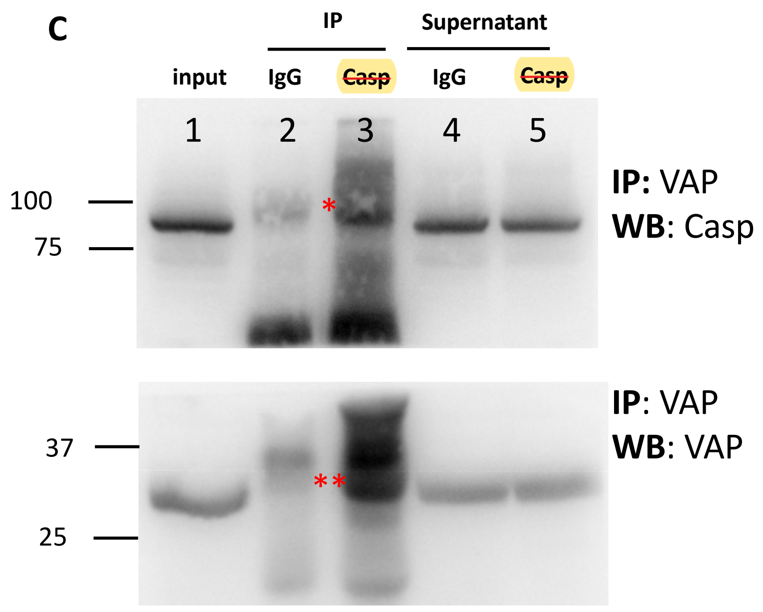


**Suppl. Figure 2. Validation of the Caspar antibody**

**A.** Western blot for different genotypes, using the anti-rabbit Caspar antibody (Materials & Methods) as a probe. Caspar cross-reacts with a ~80 kD band in adult lysates (Lane 1). When tested on animals that lack Caspar (Lane 3, Caspar$^{lof}$/Caspar$^{lof}$, or Lane 4 (Casparl$^{of}$/Caspar(Df)), the band is missing. There is also the expected increase in the intensity of the when Caspar is overexpressed using a constitute Da promoter (Lane 2).

**B.** *Caspar* was expressed and knocked down in the Dorsal compartment of the wing imaginal disc using the Apterous-Gal4 driver. Top Panel, Overexpression of Caspar:HA could be visualised using the anti-Caspar antibody, which showed the same expression pattern as the anti-HA antibody. Bottom Panel, Knockdown of *Caspar* in the Dorsal compartment led to a strong reduction in native Caspar protein expression. The ventral compartment, not affected by the knockdown, still shows normal levels of Caspar.

**C.** A affinity pulldown experiment in adult whole body lysates (See Materials & Methods) using the Rb anti-VAP antibody. VAP was successfully enriched (lane 3, bottom panel) and Caspar was enrichd (top panel, lane 3) as an interactor of VAP.

**D.** Table for peptide counts for LC-MS data from Caspar IP's of fly lysates.

**D**

Adult Lysate

| Protein | Flybase ID | igG (Peptide Count) N=4 | IP (Peptide Count) N=4 |
|---|---|---|---|
| casp | FBgn0034068 | 0 | 33 |
| Cds | FBgn0010350 | 0 | 2 |
| Gp93 | FBgn0039562 | 0 | 3.5 |
| waw | FBgn0024182 | 0 | 2.25 |
| TER94 | FBgn0286784 | 0 | 3.5 |
| CG8635 | FBgn0033317 | 0 | 3 |
| CG1234 | FBgn0037489 | 0 | 3 |
| mRpS22 | FBgn0039555 | 0 | 4 |
| Tor | FBgn0021796 | 0 | 3 |
| l(2)gl | FBgn0002121 | 0 | 3.5 |
| Naprt | FBgn0031589 | 0 | 3.5 |
| CG31694 | FBgn0051694 | 0 | 4 |
| Ppox | FBgn0020018 | 0 | 5 |
| eIF3f1 | FBgn0037270 | 0 | 3 |
| CG12237 | FBgn0031048 | 0 | 3 |
| Spn28B | FBgn0083141 | 0 | 3 |
| D2hgdh | FBgn0023507 | 0 | 3 |
| CG9253 | FBgn0032919 | 0 | 3 |
| dco | FBgn0002413 | 0 | 2 |
| CG2260 | FBgn0030000 | 0 | 2 |
| CG32409 | FBgn0052409 | 0 | 2 |

Embryonic Lysate

| Protein | Flybase ID | igG (Peptide Count) N=1 | IP (Peptide Count) N=1 |
|---|---|---|---|
| casp | FBgn0034068 | 0 | 146 |
| TER94 | FBgn0286784 | 0 | 44 |
| Vap33 | FBgn0029687 | 0 | 10 |
| Npl4 | FBgn0039348 | 0 | 8 |
| His4:CG33871 | FBgn0053871 | 0 | 6 |
| His4:CG33873 | FBgn0053873 | 0 | 6 |
| His4:CG33875 | FBgn0053875 | 0 | 6 |
| His4:CG33883 | FBgn0053883 | 0 | 6 |
| His4:CG33885 | FBgn0053885 | 0 | 6 |
| His4:CG33887 | FBgn0053887 | 0 | 6 |
| His4:CG33889 | FBgn0053889 | 0 | 6 |
| His4:CG33891 | FBgn0053891 | 0 | 6 |
| His4:CG33893 | FBgn0053893 | 0 | 6 |
| His4:CG33895 | FBgn0053895 | 0 | 6 |
| His4:CG33897 | FBgn0053897 | 0 | 6 |
| His4:CG33899 | FBgn0053899 | 0 | 6 |
| His4:CG33901 | FBgn0053901 | 0 | 6 |
| Gapdh2 | FBgn0001092 | 0 | 5 |
| Gapdh1 | FBgn0001091 | 0 | 5 |
| Ufd1-like | FBgn0036136 | 0 | 4 |
| CCT6 | FBgn0027329 | 0 | 4 |

Suppl. FIGURE 3

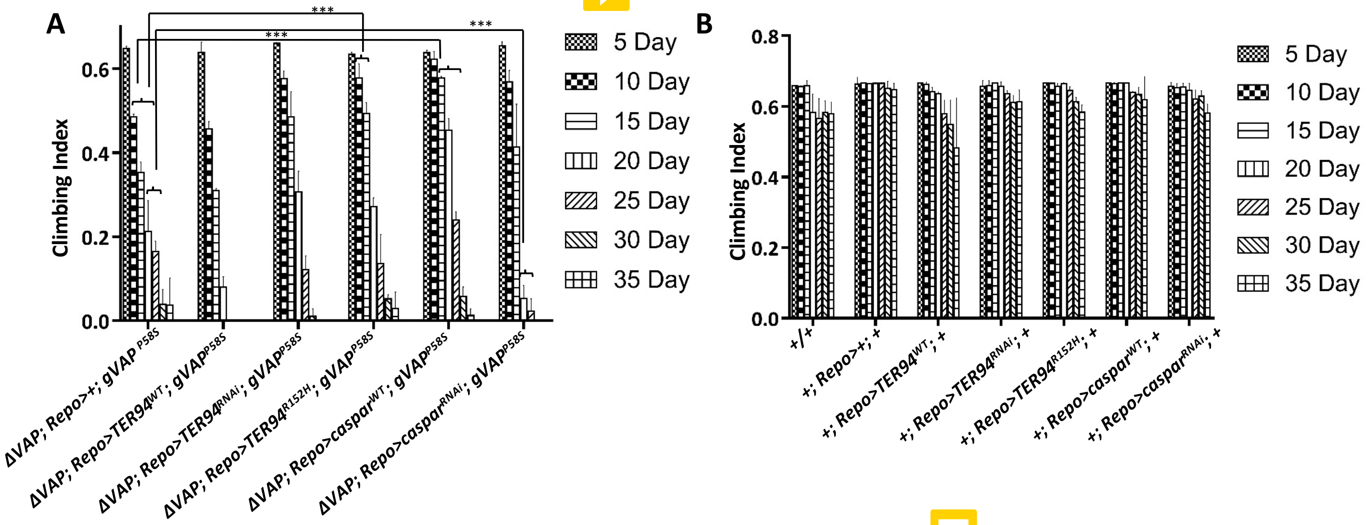


**Suppl. Figure 3. Overexpression of Caspar in the glia delays motor dysfunction.**

**A.** Climbing indices for glial overexpression of $TER94^{WT}$, $TER94^{RNAi}$, $TER94^{R152H}$, $caspar^{WT}$ and $caspar^{RNAi}$ in the $VAP^{P58S}$ background, as plotted in Fig. 3E are represented/reported in a 5-day format. p-values are as reported in Fig. 3G.

**B.** Climbing indices for glial overexpression of $TER94^{WT}$, $TER94^{RNAi}$, $TER94^{R152H}$, $caspar^{WT}$ and $caspar^{RNAi}$ in a wild-type background, as plotted in Fig. 3E are represented/reported in a 5-day format. p-values are as reported in Fig. 3H.

SUPPL. FIGURE 4

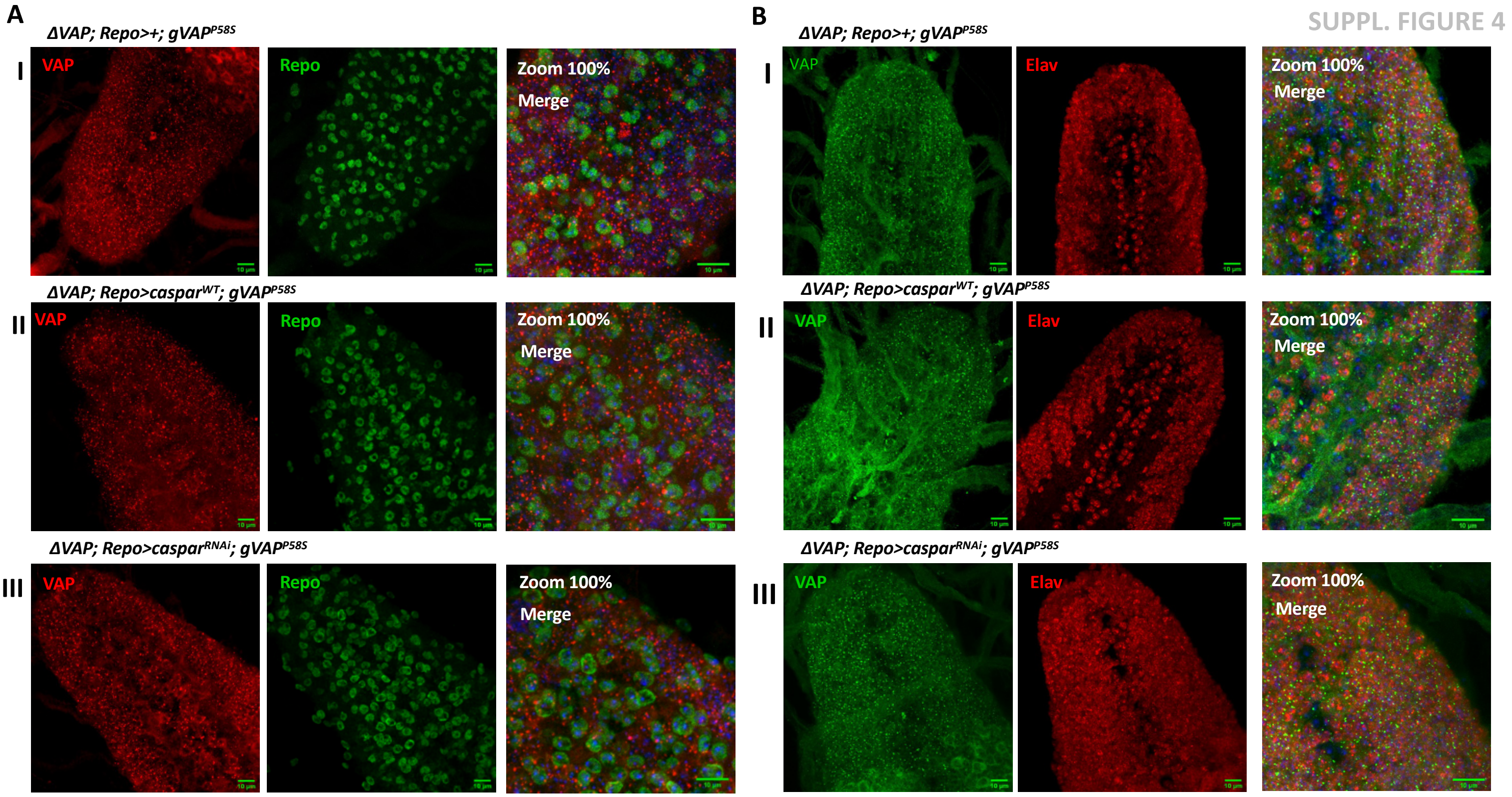


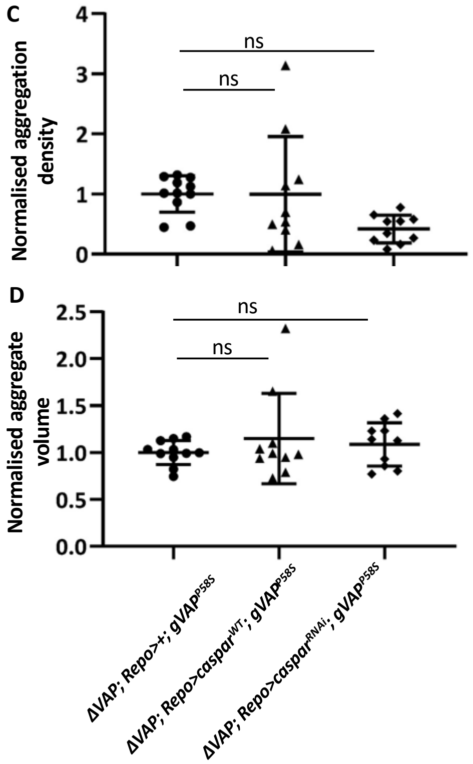


**Suppl. Figure 4. VAP inclusions in the larval brain are not significantly affected by Caspar OE.**

A. Representative images of VAP inclusions in third instar larval brains of *ΔVAP; Repo>+; gVAP*$^{P58S}$ (I), *ΔVAP; Repo>caspar*$^{WT}$*; gVAP*$^{P58S}$ (II) and *ΔVAP; Repo>caspar*$^{RNAi}$*; gVAP*$^{P58S}$ (III). The markers used are VAP, Repo and DAPI. Third panel represents merged image at 100% zoom.

B. Representative images of VAP inclusions in third instar larval brains of *ΔVAP; Repo>+; gVAP*$^{P58S}$ (I), *ΔVAP; Repo>caspar*$^{WT}$*; gVAP*$^{P58S}$ (II) and *ΔVAP; Repo>caspar*$^{RNAi}$*; gVAP*$^{P58S}$ (III). The markers used are VAP, Elav and DAPI. Third panel indicates merged image at 100% zoom.

C. Graphical representation of normalized aggregation density and normalized aggregate volume. The genotypes compared are *ΔVAP; Repo>+; gVAP*$^{P58S}$ */+* (control), *ΔVAP; Repo>caspar*$^{WT}$*; gVAP*$^{P58S}$, and *ΔVAP; Repo>caspar*$^{RNAi}$*; gVAP*$^{P58S}$. n= 10 brain samples. One-way ANOVA followed by Tukey's multiple comparison (*P<0.05, ***P<0.001, ****P<0.0001; ns, not significant). Error bars indicate s.d.

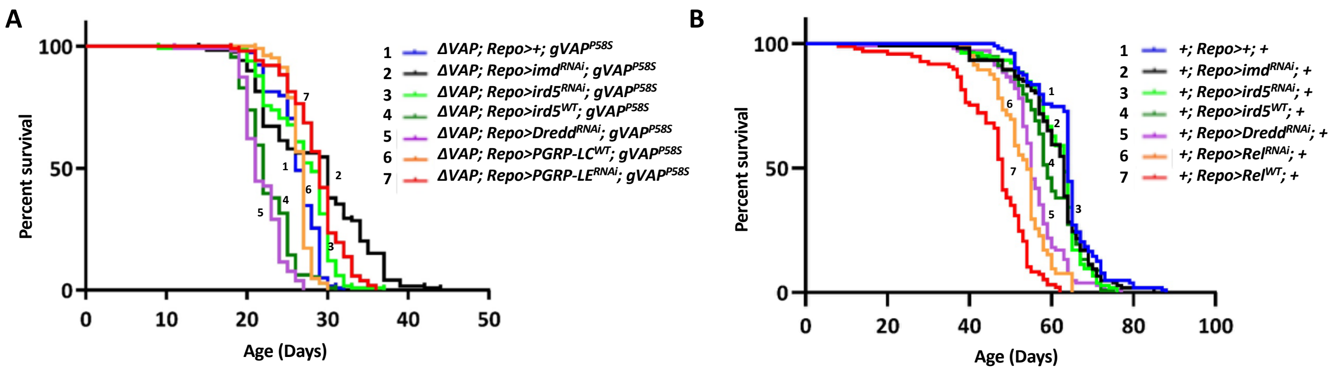


**Suppl. Figure 5. Inflammation in glia, regulated by IMD/REL signalling contributes to the progression of the disease.**

**A.** Lifespan curves for overexpression of *Imd$^{RNAi}$* (2, black curve), *ird5$^{RNAi}$* (3, light green), *ird5$^{WT}$*(4, dark green) *Dredd$^{RNAi}$* (5, purple), *PGRP-LC$^{WT}$* (6, orange) and *PGRP-LE$^{RNAi}$* (7, red) in the *ΔVAP; Repo-Gal4; gVAP$^{P58S}$* background. *ΔVAP; Repo>+; gVAP$^{P58S}$/+* (1, blue, Median= 26 Days) was used as the control. Curve comparison was done using log-rank (Mantel-Cox) test. Combined *p*-value for the whole set is <0.0001.

**B.** Lifespan curves for overexpression of *Imd$^{RNAi}$* (2, black), ird5$^{RNAi}$ (3, light green), *ird5$^{WT}$*(4, dark green) *Dredd$^{RNAi}$* (5, purple), *Rel$^{RNAi}$* (6, orange) and *Rel$^{WT}$* (7, red) in the *+; Repo-Gal4; +* background. *+; Repo>+; +*(curve 1 in blue, Median= 64 Days) was used as the control. Curve comparison was done using log-rank (Mantel-Cox) test. Combined *p*-value for the whole set is <0.0001.
